# Common, species-specific, and accession-specific responses of foliar phytohormones and morphological traits to drought and herbivory

**DOI:** 10.64898/2026.03.30.715323

**Authors:** Xue Xiao, Kruthika S. Aragam, Andrea Bräutigam, Thomas Dussarrat, Selina Gaar, Maximilian Hanusch, Robin Heinen, Marvin Hildebrandt, Ruth Jakobs, Robert R. Junker, Radhika Keshan, Judit V. Mendoza Servín, Elikplim A. Setordjie, Yonca B. Seymen, Anke Steppuhn, Sybille B. Unsicker, Nicole M. van Dam, Baris Weber, Sarah K. Weirauch, Wolfgang W. Weisser, Dominik Ziaja, Jörg-Peter Schnitzler, J. Barbro Winkler, Caroline Müller

## Abstract

**Background:** Plants are exposed to various environmental challenges. With ongoing climate change, droughts and insect outbreaks are expected to become more frequent. Thus, a better understanding is needed of how different plant species respond to such single and combined challenges. This study investigated common versus species-specific responses to environmental challenges in three perennial plant species of different growth forms and whether responses differ intraspecifically among accessions. Clones of different accessions of the herbaceous species *Tanacetum vulgare*, the woody vine *Solanum dulcamara*, and the tree *Populus nigra* were subjected to similar control, herbivory, drought, and combined (drought and herbivory) treatments for the same periods. After the exposure, concentrations of foliar phytohormones and various morphological traits were measured.

**Results:** Across all species, several foliar phytohormones and one of ten morphological traits responded consistently to the environmental challenges. Jasmonoyl-isoleucine was induced by herbivory and the combined treatment, abscisic acid (ABA) by drought and the combined treatment, and indole acetic acid by the combined treatment in all species. Root mass remained unchanged in all species. However, structural equation models (SEMs) revealed a shared regulatory pathway across species in which ABA connected treatment and root mass, indicating a common hormonal response potentially linking challenges to growth responses. Despite these common patterns, species-specific responses were pronounced. In *P. nigra*, a unique induction of salicylic acid was found under the combined treatment, while aboveground mass and root-shoot ratio remained unaffected by any treatment, in contrast to the other two species. Species-specific SEMs further indicated distinct phytohormone-mediated pathways underlying morphological variation. Phenotypic plasticity reflected these species-specific patterns, with none of the phytohormones or morphological traits exhibiting uniform plasticity across species. Intraspecific variation further shaped responses, as phytohormone and morphological trait plasticity depended on accession, indicating substantial accession-specific plant responses.

**Conclusions:** Our results indicate that some responses to comparable challenges may be conserved across species, while others are species-specific. The combined treatment elicited the most pronounced responses, and such complex responses may become more frequent under current global change. Our study highlights that comprehensive understanding of plant responses requires systematic comparisons at both interspecific and intraspecific scales.

## Background

Plants are constantly exposed to diverse abiotic and biotic challenges. Among these, drought and insect herbivory are particularly important because both are predicted to occur more frequently under current climate change scenarios and can strongly impair plant performance, growth, and survival [1–3]. Plant responses to such challenges are largely regulated by phytohormones, which integrate environmental signals with developmental programs, and thereby shape physiological and morphological adjustments [4]. Thus, phytohormone signaling as an early molecular response to environmental challenges enables adjustments of growth and development and coordinated patterns of resource allocation [5]. The magnitude of these responses depends on the phenotypic plasticity of the individuals, that is, their ability to modify trait expression in response to environmental conditions [6]. Responses to abiotic or biotic challenges have often been studied in isolation, although they commonly co-occur in the environment [7, 8]. Phytohormone signaling can help plants to alter their phenotypes also under exposure to such co-occurring challenges. However, comparative studies that expose plants from different families with distinct growth forms to comparable challenges are lacking, but would enable to disentangle common versus species-specific responses as well as intraspecific variation in plasticity. In response to drought, especially abscisic acid (ABA) is a key regulator as it directly promotes stomatal closure and thus prevents water loss [9]. In addition, ABA signaling alters gene expression entailing various physiological responses, which impact root growth [10]. The phytohormone indole-3-acetic acid (IAA) is mainly involved in cell elongation and root growth, but also linked to drought tolerance in plants [11], particularly when interacting with elevated ABA concentrations [12]. Both ABA and IAA have been found to increase or decrease in response to drought, depending on the plant species and the drought intensity [9]. In addition, salicylic acid (SA) promotes osmotic regulation under water deficit by enhancing levels of soluble sugars and proline [13, 14]. Jasmonic acid (JA) and its bioactive derivative jasmonoyl-isoleucine (JA-Ile) have also been found to increase in response to drought [9]. JA, JA-Ile, and SA can interact positively with ABA signaling to induce stomatal closure [15, 16]. Thus, drought induces a coordinated regulatory network integrating various phytohormone signaling pathways [4, 9]. In addition to these signaling responses, drought also directly reshapes plant morphology, with relatively more investment into root compared to shoot biomass, resulting in an enhanced root to shoot ratio [17].

Beyond drought, phytohormones regulate plant responses to herbivory, in which the JA pathway plays a central role [4]. The levels of JA and JA-Ile usually increase when insects with chewing biting mouthparts damage the tissue, leading to the induction of direct and indirect defenses [18]. In concert with JA, the phytohormones SA, ABA, and others, such as ethylene (ET), are responding, but in a highly challenge-specific way [19]. The SA pathway sometimes antagonizes JA/ET signaling [20]. Tissue damage imposed by herbivores together with responses mediated by phytohormone signaling suppress plant growth, partly caused by trade-offs in resource allocation between induced defense and growth [18]. Since the same phytohormones are involved in mediating plant responses to both drought and herbivory, and due to crosstalk among the hormones, exposure to combined challenges may modulate the resulting morphological responses. Indeed, drought (seven days) combined with feeding by a specialist herbivore enhanced JA and SA levels but suppressed ET production in *Solanum dulcamara* (Solanaceae), while drought alone mostly enhanced ABA, with levels not increasing when herbivory was taking place in combination [21], indicating that effects are not simply additive. In *Populus nigra* (Salicaceae), herbivory by leaf-chewing insects led to a pronounced induction of ABA and JA, but not SA, while drought alone induced ABA [22, 23]. Aboveground dry mass of *Tanacetum vulgare* (Asteraceae) decreased under drought (three weeks) and under combined challenges of drought and herbivory by the generalist *Spodoptera exigua* (five days), whereas herbivory alone had no effect [8].

Responsiveness may not only differ among species, but also intraspecifically. For example, genotype-specific responses of JA, JA-Ile, SA, ABA, and IAA to galling herbivores were found in *Populus angustifolia* [24]. In *T. vulgare*, plants of several accessions (maternal origins) differed in their plasticity of leaf terpenoid richness under environmental challenges [8]. In *P. nigra*, differences in growth patterns were also found between female and male trees; for example, females grew taller than males in a riparian forest [25]. However, such individual findings do not allow for general conclusions on plasticity, but experiments are needed that apply comparable treatments of single and combined challenges across plant species and accessions or genotypes.

In this study, we investigated general, species-specific, and accession-specific responses in phytohormones and morphological traits of three perennial plant species from different plant families and with different growth forms to single and combined challenges. *Tanacetum vulgare* was chosen as an herbaceous perennial, *Solanum dulcamara* as a perennial woody vine, and *Populus nigra* as a long-lived, dioecious tree. Within each species, several accessions were clonally propagated, and the resulting clones were exposed to either no challenge as control, drought, herbivory, or the combination of drought and herbivory under standardized conditions to assess the phenotypic plasticity of the three plant species. At the end of the exposure time, we determined the concentrations of five foliar phytohormones and ten plant morphological traits. We hypothesized that ABA and IAA are enhanced under drought, while JA and JA-Ile are induced under herbivory in all three species. Growth-related traits, such as aboveground mass, were expected to be reduced under both drought and herbivory alone, with a stronger reduction under the combined challenge, while root–shoot ratio should increase in all species. However, we expected that the magnitude of phytohormonal responses and morphological phenotypic plasticity would differ across species. Finally, we explored the relationships between phytohormones and morphological traits across species and accessions to deliver potential mechanistic explanations.

## Materials and methods

### Plant and insect rearing

Plants of the herb *Tanacetum vulgare*, the vine *Solanum dulcamara*, and the tree species *Populus nigra* were collected in the field (without special collection permission required) and reared in different research groups. To test common, species-specific, and accession-specific responses of these plants to comparable challenges, experiments were performed under highly controlled environmental conditions in phytotron chambers of the ExpoSCREEN facility at Helmholtz Munich [26]. Due to space limitation in the climate chambers, species were examined sequentially (for time schedule see Table S1).

Seed heads of six different accessions of *T. vulgare* were collected near Bielefeld, Germany, and a stock of offspring plants, representing different terpenoid leaf chemotypes, established at Bielefeld University (for details see [27]and Table S2). Stock plants were shipped to Munich in a steamed 1:1 mixture of sand: soil (Fruhstorfer Einheitserde Typ T, Gebr. Patzer GmbH & Co. KG, Sinntal-Altengronau Germany). Per accession-chemotype combination (total of 12), 16 clones were prepared from root and stem cuttings. From *S. dulcamara*, three accessions were collected in Germany and maintained at Hohenheim University (Stuttgart, Germany) and three accessions were collected in the Netherlands and subsequently maintained as clonal stocks at Leipzig University [28] (for details, see Table S3). Per accession, 32 clones were propagated by stem cuttings and sent to Munich. Trees (accessions) of three males and three females of *P. nigra* were originally obtained from a tree nursery (Waller Baumschulen, Schwäbisch Hall, Germany; for details, see Table S4). Per accession, 16 stem cuttings were taken from young trees grown in pots in a greenhouse at Kiel University. Cuttings were rooted in stone wool (Grodan, Roermond, The Netherlands) and subsequently transplanted into 7 cm x 7 cm x 8 cm pots filled with lightly fertilized growing substrate (seedling substrate; Klasman-Deilmann, Geeste, Germany) prior to shipping to Munich.

After arrival in Munich, plants were acclimatized over a period of time appropriate to the developmental stage of each species (Table S1). All plants were then transferred to an 8: 1: 1 mixture of peat substrate (Einheitserde CLT, Balster Einheitserdewerk, Fröndenberg, Germany): sand (0.6 – 1.2 mm): vermiculite in 13 x 13 x 13 cm pots and grown in a greenhouse at 16 h:8 h (day/night), before they were placed into the phytotron chambers at least a week before the experimental treatments for acclimatization to the growth conditions (for detailed facility parameters, see [29]. Climatic conditions in the phytotron chambers simulated average June conditions in Northern Germany, with day/night temperatures of 25°C/14.5°C, relative humidity of 40%/80%, and a 16 h photoperiod. To simulate the natural diurnal light cycle, light intensity was ramped up over 3.5 h to 718 μmol m^−2^ s^−1^ photosynthetic photon flux density (PPFD) at canopy level, maintained at this level for 9 h, and reduced over the following 3.5 h to darkness. To enable standardized sampling, treatments were started in two out of four chambers one day apart (Fig. 1), and the diurnal program was shifted by 1 h between chambers harvested on the same day.

**Fig. 1.**
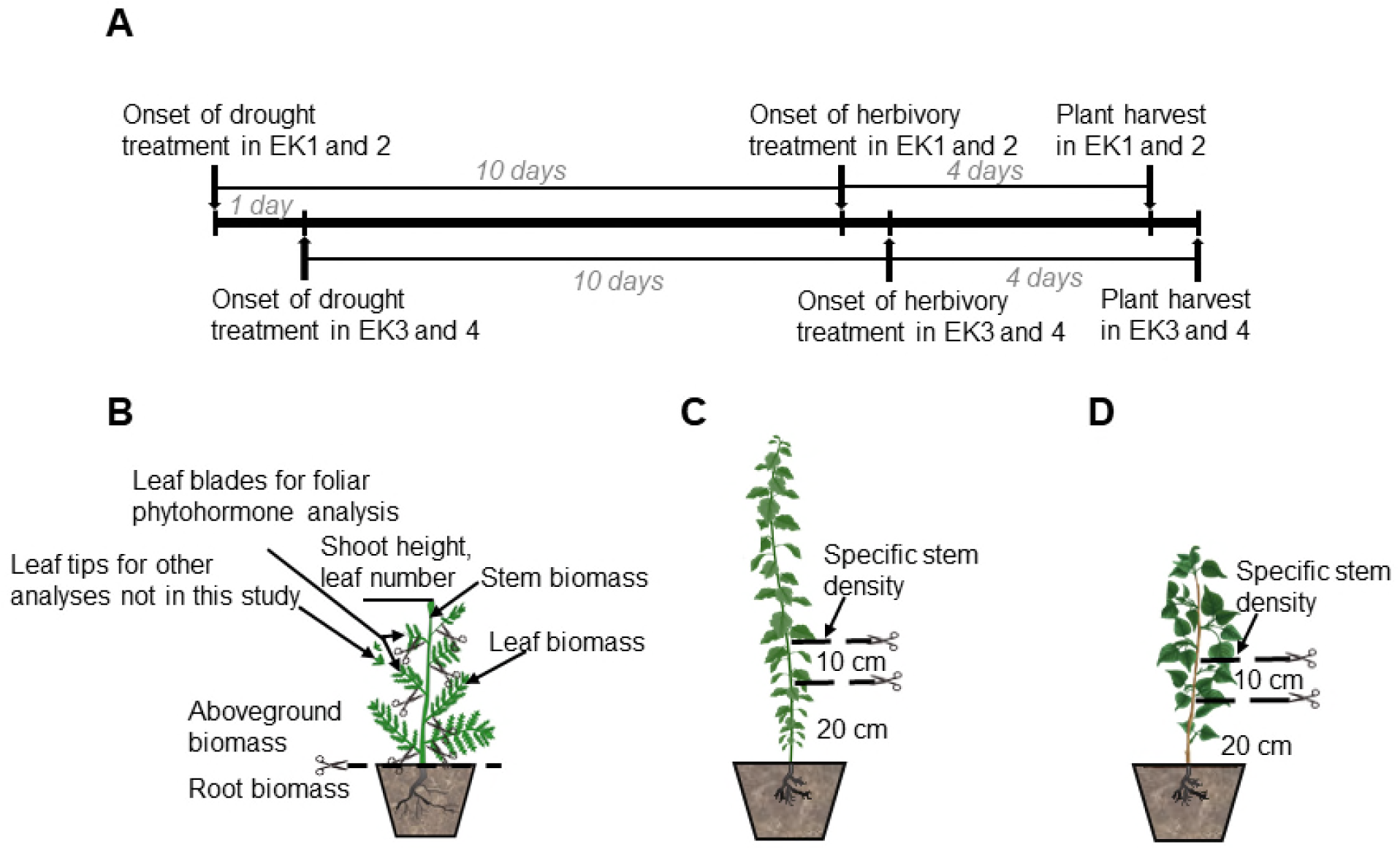
Experimental timeline and plant harvest design for the three experimental species. The timeline was identical for all species (A). Morphological traits assessed at harvest were consistent across species, except specific stem density, which was not measured in *Tanacetum vulgare* (B), but measured in *Solanum dulcamara* (C) and *Populus nigra* (D). EK – experimental chamber.

Larvae of *Spodoptera exigua* were used as herbivores for *T. vulgare* and *S. dulcamara*. They were reared at Hohenheim University on a bean flour-based artificial diet (as described in [30]) and maintained on this diet in ventilated boxes in a climate cabinet at 22°C, 16:8 light: darkness (L:D), and 60% relative humidity (RH) until shipped to Munich three days prior to the start of the experimental herbivore exposure. In Munich, larvae were kept in the greenhouse (22°C, 16:8 L:D, 60% RH) and fed with leaves of the same accessions used in the respective experiment with *T. vulgare* or *S. dulcamara.* As *S. exigua* did not feed on *P. nigra* during preliminary trials, larvae of the feeding generalist *Lymantria dispar* were used for the *P. nigra* experiment instead. Larvae of *L. dispar* hatched from eggs obtained from the US Department of Agriculture (Buzzards Bay, MA, USA) without permissions needed and were reared on artificial diet (MP Biomedicals LLC, Illkirch, France) under room temperature in the laboratory. Four days before the onset of the experiment, larvae were shipped to Munich and conditioned to feeding on *P. nigra* by supplementing their diet with a mix of leaves from the tree accessions used in the experiment.

### Experimental set-up

This experiment included additional measurements that are not addressed in the present manuscript. Here, we focus on phytohormones and morphological traits, while other aspects of the experiment will be explored in subsequent studies.

For each plant species, clones of the same accession of comparable size were assigned to one of the four treatments: control, drought, herbivory, and combined treatment (drought and herbivory) (see Table S5A-C for water regimes of each species). One (*T. vulgare* or *P. nigra*) or two clones (*S. dulcamara*) per accession were placed in each of four sub-chambers within each of the four main phytotron chambers (Figs S1-3). In each main chamber, one of the four treatments was assigned to one sub-chamber, preventing communication through the air between plants exposed to different treatments. Shortly before plants were transferred into the sub-chambers, a polyethylene terephtalate (PET) bag (RUBIN; Dirk Rossmann GmBH, Burgwedel, Germany) was fixed at the rim of each pot and pulled up later, when the insect herbivory treatment started. Four days before the onset of the drought treatment, pictures of every plant were taken from different angles in a photostation, as described in [29] to estimate leaf area (data not presented here). Drought conditions in both the drought and combined treatments were initiated by withholding one irrigation event and subsequently providing only 50% of the water supplied to the other two treatments (control, herbivory) until the end of the experiment (see Table S5A-C for watering amounts and Table S6A-C for soil water content). In cases of severe wilting, 50 to 200 mL of water were supplied in addition to the automatic irrigation, depending on the treatment and species (Table S5). Ten days after initiation of the drought treatment, the herbivory treatment was initiated by placing groups of three larvae (two 3^rd^ instar and one 4^th^ instar) of either *S. exigua* or *L. dispar* on each of the experimental plants of the herbivory and combined treatments (Fig. 1). To standardize the treatments across plant species, larvae were weighed as groups, before being placed on the plants one hour after onset of light. The PET bags previously kept at each pot rim were carefully lifted and kept open at the top to prevent larval escape while maintaining airflow. The bags exceeded the height of the plants of each treatment by approximately 30 cm. Three days later, the roasting bags were closed at the top with paper clips to reduce airflow for collection of volatile organic compounds (data not presented here). One day later, the roasting bags were removed from all plants, larvae were collected, and the final mass of the groups of (remaining) larvae per plant was recorded to evaluate their average growth rate.

### Plant harvest

Immediately after herbivore removal, leaf tips (∼ 1cm) were collected from at least three leaves per plant for subsequent RNA sequencing and microbiome analyses (data not shown). The corresponding leaf blades were harvested within 5 minutes, wrapped in aluminum foil, and frozen in liquid nitrogen for phytohormone analysis. The remaining plant was then screened in the photo station as described above. The leaf number was counted and plant height was measured. Leaves were collected and scanned with a flat-bed scanner (Epson Perfection V500 Photo, Epson, Düsseldorf, Germany) (except for *T. vulgare*, for which scanning was not feasible due to the pinnatisect leaves that curled severely under drought and herbivory treatments). Leaves were weighed fresh and again after drying at 70°C for three days. Leaf water content was calculated as (fresh mass-dry mass)/dry mass. A 10-cm stem segment was cut 20 cm above the soil surface, and its basal diameter and fresh mass were measured, expect for *T. vulgare* because not all plants developed stems in that species. Stem segments were subsequently dried (as for leaves) and reweighed to calculate specific stem density (stem dry mass/ fresh volume). The remaining stem material was harvested, dried under the same conditions as leaves and stems, and weighed. Total aboveground dry mass was determined from the sum of all dried leaf and stem material that was weighed separately. All pots were stored at 4°C until roots were washed free of soil, which was completed within two days of sampling. Cleaned roots were then dried (as above) and weighed.

### Phytohormone analysis

The flash-frozen leaf blades were lyophilized in aluminum wraps for 48 hours with a freeze-dryer (Alpha 1-4 LSC, Martin Christ) and homogenized in a mill (Retsch MM400, Retsch GmbH, Haan, Germany). An aliquot of 15 mg (+/- 1) of the dried leaf material was shipped to University of Hohenheim and stored at −20°C until extraction for analyses of JA, JA-Ile, SA, ABA, and IAA. This leaf aliquot was homogenized in 1 mL ethyl acetate spiked with deuterated phytohormones as internal standards [60.4 ng D6-JA (HPC Standards GmbH, Cunnersdorf, Germany), 14.2 ng D6-JA-Ile (HPC Standards GmbH, Cunnersdorf, Germany), 20 ng of each of D4-SA, D6-ABA (OlChemIm Ltd., Olomouc, Czech Republic), and D5-IAA (Hölzel Diagnostika Handels GmbH, Köln, Germany)] with 6-7 ceramic beads (Zirconox®) in a Bead Mill 24 (Fisherbrand^TM^) twice for 20 sec at 4 m/s. After centrifugation at 4°C, the pellet was re-extracted for 15 sec at 4 m/s in 1 mL ethyl acetate and centrifuged again. The combined supernatants were dried in a vacuum concentrator (Eppendorf, Hamburg, Germany) and reconstituted in 400 µL 70% methanol (v/v) with 0.1% formic acid by vortexing for 10 min. The extracts were stored at −20°C, until 4 µL were subjected to analysis by UPLC-ESI-MS/MS SCIEX QTRAP 6500+ (Agilent 1290 II, Agilent Technologies, USA). Phytohormones were separated on a C18 column (Waters UPLC column ACQUITY PREMIER PST HSS T3 C18, 100 Å, 1.8 µm, 2.1x 150 mm, Milford, MA, USA) at 40°C with a flow rate of 0.35 mL/min with water and methanol (both containing 0.2% formic acid) as solvents A and B in gradient mode (solvent B: 0 min: 35%; 2 min: 35%; 4 min: 38%; 6 min: 65%; 8 min: 75%; 8.2 min: 100%; 9.5 min: 35%). All except one phytohormone were detected in negative ionization mode with parent/daughter ion selections of 209/59 (JA), 215/59 (D6-JA), 322/130 (JA-Ile), 328/130 (D6-JA-Ile), 263/153 (ABA), 269/159 (D6-ABA), 137/93 (SA), 141/97 (D4SA), while IAA (176/30) and D5-IAA (181/134) were detected in positive ionization mode. Upon peak area integration, phytohormones were quantified relative to the internal standards.

### Statistical analyses

All statistical analyses were conducted in R v4.5.0 [31], using the packages dplyr [32], tidyverse [33], janitor [34], nlme [35], DHARMa [34], glmmTMB [36], MuMin [37], car [38], flextable [39], officer [40], multcomp [41], multcompView [42], vegan [43], piecewiseSEM [44,45], and semEff [45]. Visualization was done with ggplot2 [46] and patchwork [47].

We first estimated the effects of species on each phytohormone and morphological trait based on datasets excluding missing values of corresponding response variables, such as specific stem density in *T. vulgare*. Following that, the effect of treatment and accession on each phytohormone and morphological trait was tested within each species, accounting for the slightly different data structure per species, using linear models (LMs). In addition, sex was included in models of *P. nigra*. Because accession and sex were collinear in *P. nigra* (three accessions female, three accessions male), the interaction between accession and sex was not considered. Response variables were transformed either via square root or log-transformed when this improved the model performance based on DHARMa Q-Q plots. We then evaluated the significance of predictors in the most parsimonious models (the model with the least degree of freedoms with ΔAICc > 2) using Type II ANOVA. For *T. vulgare* and *S. dulcamara*, both treatment and accession were included in the final models as no multicollinearity was detected between these predictors.

To assess the effect of drought on herbivore performance within each species, we fitted linear models using relative average herbivore growth rate as the response variable. Treatment (here: herbivory vs. combined) and the initial average herbivore mass prior to application to plants were included as explanatory variables. Treatment effects, independent of initial herbivore biomass, were evaluated using Type II ANOVA.

We quantified phenotypic plasticity of each trait between clones of a given plant growing in the sub-chambers of the same main chamber as previously described in[48], employing the relative distance plasticity index (RDPI) = (|x_c_ – x_s_|)/(|x_c_ + x_s_|), where X_c_ and X_s_ represent the different trait values of clones kept under control (c) and challenge (s) conditions. The RPDI value ranges from 0 (no plasticity) to 1 (maximal plasticity). Data of clones of the same accession were excluded if any trait values were missing from one of the four sub-chambers at comparable position, resulting in final data sizes of 148 plants for *T. vulgare*, 184 plants for *S. dulcamara*, and 96 plants for *P. nigra*. We then analyzed the effects of treatment, accession, and sex in the case of *P. nigra*, using generalized linear effect models implemented in glmmTMB with a beta error distribution and logit link function, appropriate for continuous proportion data bounded between 0 and 1 that exhibit strong right-skewness. We evaluated predictor significance in the most parsimonious models (the model with the least degree of freedoms with ΔAICc > 2) using Type II ANOVA.

To illustrate sample similarity and assess effects of species and treatment on variation in phytohormone profiles and morphological traits, we performed a joint Redundancy Analysis (RDA) using phytohormone profiles and morphological traits as response variables and species and treatment as explanatory variables across all three species. Prior to analysis, phytohormone concentrations were normalized to the median across plants of all species and log-transformed to reduce skewness. The resulting phytohormone values as well as the raw morphological trait data were z-transformed across species, ensuring comparability for multivariate analyses. In addition, separate RDAs were conducted for each species using similar processing. Traits (phytohormones and morphological traits) that were most strongly associated with species separation in the RDA (as indicated by longer vectors) and were significantly affected by treatment in at least one species were visualized using box plots.

To investigate the relative influence of treatment and accession on phytohormone and morphological responses, and to quantify the potential mediating role of phytohormones, we fitted piecewise structural equation models (SEMs). Before the analysis, phytohormone concentrations and morphological traits of each species were processed following the same procedure as for the RDA. SEMs were fitted using LMs. For each response variable, the model was constructed based on *a priori* knowledge of the study systems (Fig. S4). During model validation, missing paths (i.e., previously unconsidered significant relationships) were evaluated and included if biologically justified, or otherwise left to freely covary. Potentially missing paths were evaluated with the d-separation test and global model fit was evaluated via Fisher’s C (*p* > 0.05). Bootstrapped standardized effects were computed on each SEM using 1,000 iterations to extract direct, indirect, mediated, and total effects. SEM plots present direct paths from treatments and phytohormones to morphological traits. Absolute mediator sizes of phytohormones between treatment or accession and morphological traits were also plotted. Within *P. nigra*, we compared the effects of treatment in combination of either accession or sex. For each set of SEMs, we calculated the sum of Akaike weights as an indicator of model performance. The SEM exhibiting substantially better performance (i.e., lower AIC with ΔAICc > 2) is presented in the main text. Because this study focused on the direct effects of treatment and assessment on phytohormones and morphological traits, indirect links among phytohormones were not included in the structural equation models (SEMs).

## Results

### A subset of phytohormone responses to treatments are species-specific

All foliar phytohormone concentrations differed significantly among species (Table 1) and therefore contributed to species separation (Fig. 2A; *F* = 173.60, *p* < 0.001). Leaves of *Tanacetum vulgare* tended to exhibit higher concentrations of JA, JA-Ile, ABA, and IAA than *Solanum dulcamara*, whereas *Populus nigra* had higher SA concentrations (Fig. 2A). Across all three species, concentrations of JA-Ile, ABA, and IAA were significantly affected by treatment (Table 1). JA-Ile was consistently induced by herbivory and the combined treatment (Fig. 3A–C), ABA was consistently induced by drought and the combined treatment (Fig. 3Dw -F), and IAA was consistently elevated under the combined treatment relative to controls (Fig. S5D–F). Generally, phytohormone responses were similar responses across species (Figure 2A), but detailed analyses of individual phytohormones revealed pronounced species-specific response patterns (Table 1).

**Fig. 2.**
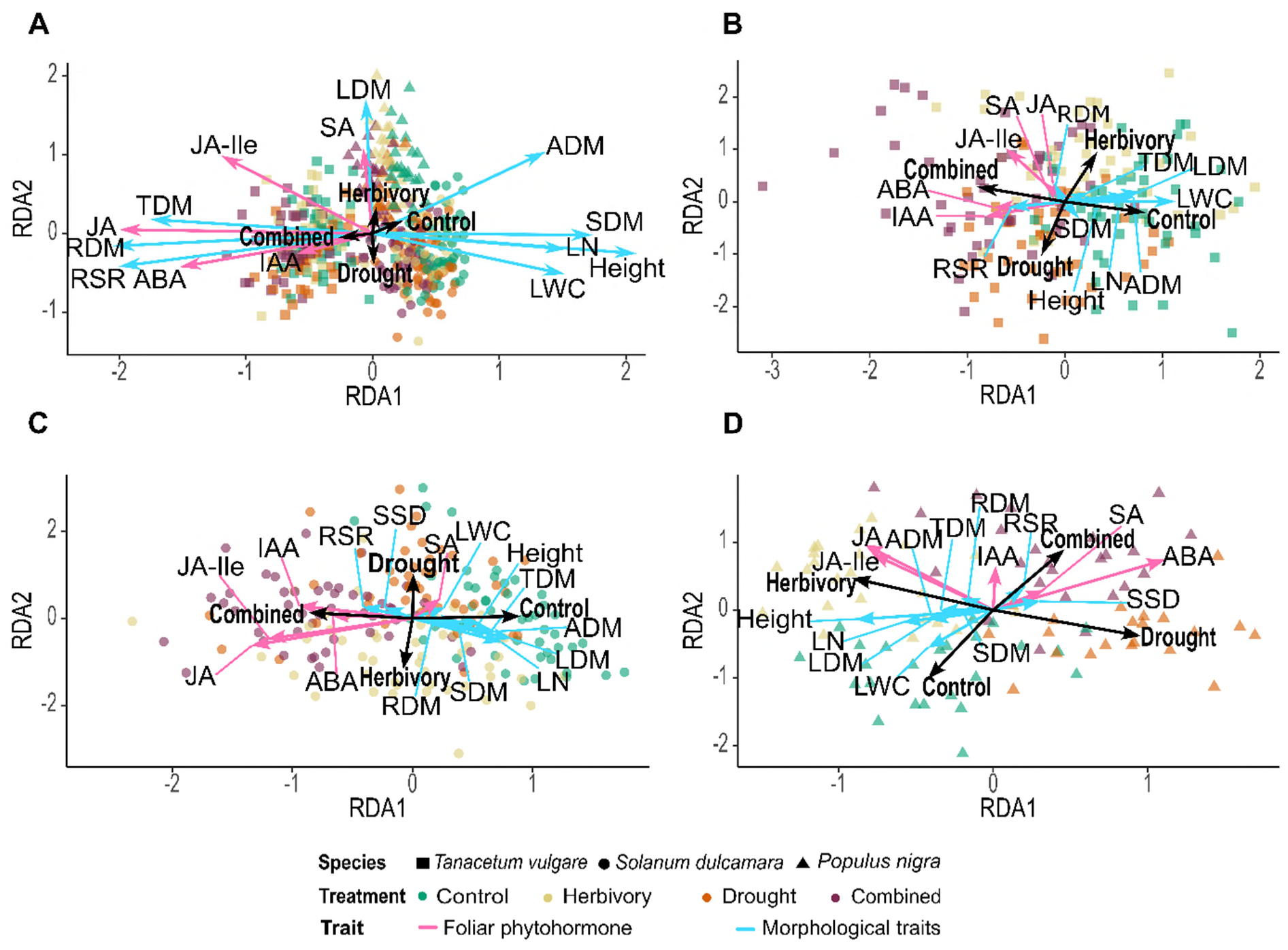
Redundancy analysis of foliar phytohormone concentrations and morphological trait across all three species (A), and separately for *Tanacetum vulgare* (B), *Solanum dulcamara* (C), and *Populus nigra* (D). The morphological traits are: LN: leaf number; LDM: leaf dry mass; SDM: stem dry mass; ADM: aboveground dry mass, RDM: root dry mass; TDM: total dry mass, RSR: root-shoot ratio; LWC: leaf water content; SSD: specific stem density. Point shapes indicate species and colors indicate treatments. Foliar phytohormones are shown with pink arrows, and morphological traits with blue arrows.

**Fig. 3.**
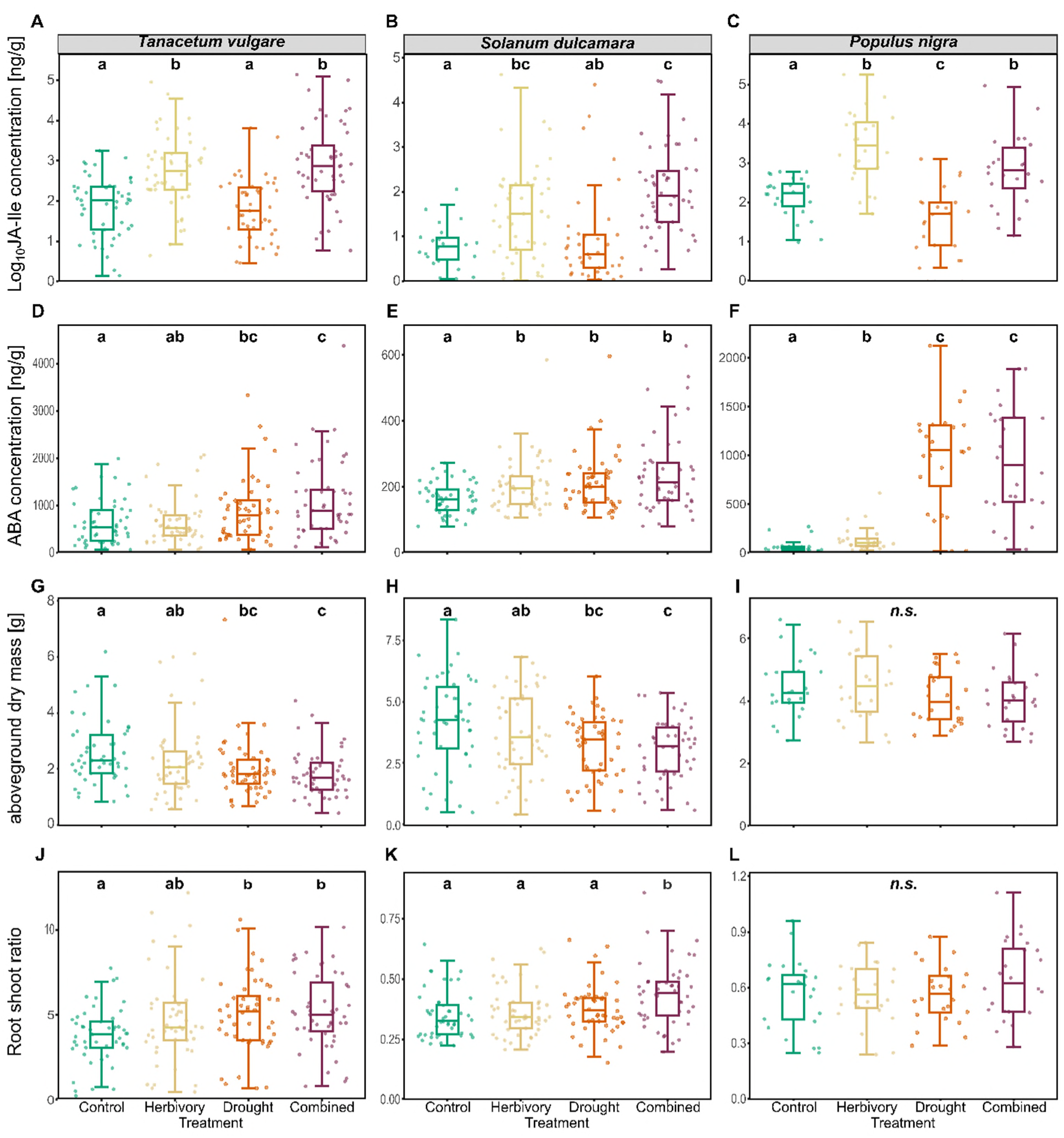
Boxplots of foliar phytohormone concentrations (JA-Ile: jasmonoyl isoleucine; ABA: abscisic acid) and morphological traits across treatments of each species, *Tanacetum vulgare* (A, D, G, J), *Solanum dulcamara* (B, E, H, K), and *Populus nigra* (C, F, I, L). Data are presented as boxplots, with medians, interquartile ranges (IQR, boxes), and whiskers extending to the most extreme values with max. 1.5 times the IQR. Individual values are plotted as dots; *n* = 48 per treatment for *T. vulgare* and *S. dulcamara*, n = 24 per treatment for *P. nigra*. Different letters indicate statistically significant differences (F-test, *p* < 0.05); *n.s.*: not significant. Panels A–F show phytohormone concentrations, with the y-axis representing concentration in ng per g of dry mass. Please note that y-axes scales differ among panels.

**Table 1.**
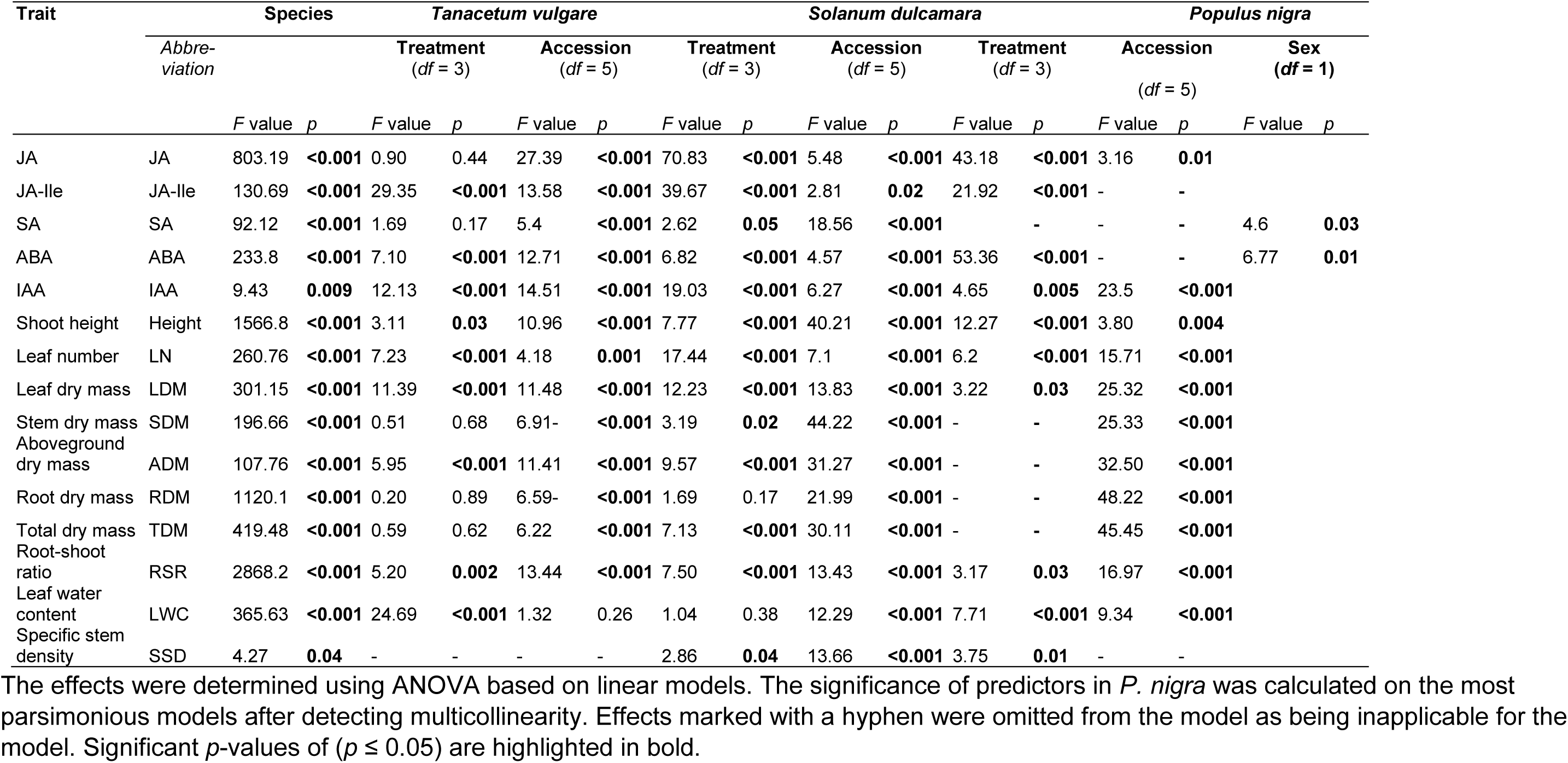
Summary of *F* and *p* values from ANOVAs testing the effects of treatment and accession on foliar phytohormone concentrations and morphological traits of *Tanacetum vulgare*, *Solanum dulcamara*, and *Populus nigra*.

In *T. vulgare*, sample separation was largely driven by treatment effects, particularly by differences between the control and the combined treatment (Fig. 2B; *F* = 5.47, *p* < 0.001). All phytohormones were positively associated with the combined treatment, most notably JA-Ile, ABA, and IAA. However, JA and SA concentrations were not significantly affected by treatment (Table 1; Figs S5A, S5G). In contrast to the other two species, ABA was not induced by herbivory in *T. vulgare*, whereas IAA increased not only under the combined treatment but also under drought (Figs 3D, S5D).

Similarly, in *S. dulcamara*, sample separation was mainly explained by differences between the control and the combined treatment (Fig. 2C; *F* = 9.25, *p* < 0.001). Concentrations of ABA and IAA, and particularly of JA and JA-Ile, were higher in the combined treatment, whereas SA was weakly increased in the control and the drought treatment. JA concentrations were highest under the herbivory and the combined treatment, intermediate under drought, and lowest in control plants (Fig. S5B). This pattern was not mirrored for JA-Ile, for which concentrations in drought-exposed plants did not differ significantly from those of the herbivory and the control treatment (Fig. 3B). Concentrations of SA remained unchanged across treatments (Fig. S5H), but ABA was induced not only by drought and the combined treatment but also by herbivory in *S. dulcamara* (Fig. 3E). IAA concentrations increased under both the drought and the combined treatment (Fig. S5E).

In *P. nigra*, samples were clearly separated among all four treatments (Fig. 2D; *F* = 5.70, *p* < 0.001). Levels of JA and JA-Ile were positively associated with herbivory, whereas SA, and ABA were associated with the drought and combined treatments. Among these phytohormones, JA, JA-Ile, and ABA showed stronger correlations with treatment than SA and IAA. JA concentrations were highest under herbivory, intermediate under the combined treatment, and lowest under the drought and control conditions (Fig. S5C). Notably, SA was induced by the combined treatment only in *P. nigra* (Fig. S5I). Concentrations of ABA were particularly higher under drought than under herbivory in *P. nigra*, although herbivory still resulted in higher ABA levels than in the control (Fig. 3F). Drought did not lead to increased IAA concentrations in *P. nigra* (Fig. S5F).

Patterns of phenotypic plasticity (RDPI, the relative variance of specific traits under one environmental challenge compared to the control) in phytohormone concentrations broadly mirrored these species-specific treatment responses (Tables 2-4). No phytohormone showed a consistent treatment effect on plasticity across all three species, but within species, accession-specific responses were found for some traits. In *T. vulgare*, the RDPIs of the JA, SA, and IAA concentrations varied among accessions, but none due to treatment (Table 2). In *S. dulcamara*, treatment effects were found in the RDPIs of the JA and JA-Ile concentrations, with higher RDPI under herbivory and the combined treatment, followed by drought, compared with control plants. In addition, the RDPI of the JA concentration varied among accessions, whereas the RDPI of the IAA concentration was influenced only by accession (Table 3. In *P. nigra*, the RDPIs of JA and ABA s were significantly affected by treatment. Concentrations of JA were highest under the herbivory treatment, followed by the combined treatment, and lowest in the drought treatment relative to the controls. The RDPI of the ABA concentration was higher under the drought and combined treatments than under herbivory (Table 4). The RDPI of the IAA concentration varied among accessions (Table 4), while the RDPI of the SA concentration was influenced by sex, with higher values in female than in male plants (Table S7, Fig. S6).

**Table 2.** Means (standard deviations) of phenotypic plasticity (measured as RDPI^1^) of foliar phytohormone concentration and morphological traits of *Tanacetum vulgare* of different accesions^2^ exposed to different environmental challenges (combined: drought and herbivory).

| Trait<sup>3</sup> | Treatment | M6<br>df = 6 | M9<br>df = 10 | M11<br>df = 4 | M16<br>df = 8 | M18<br>df = 4 | M22<br>df = 5 |  |
| --- | --- | --- | --- | --- | --- | --- | --- | --- |
| JA | Herbivory | 0.292 (0.088) | 0.215 (0.056) | 0.327 (0.085) | 0.171 (0.065) | 0.322 (0.069) | 0.204 (0.086) | Accession:<br>$\chi^2 = 19.63$;<br>$p = 0.001$ |
|  | Drought | 0.257 (0.054) | 0.156 (0.050) | 0.421 (0.085) | 0.213 (0.055) | 0.184 (0.044) | 0.258 (0.060) |  |
|  | Combined | 0.379 (0.101) | 0.156 (0.031) | 0.287 (0.073) | 0.178 (0.059) | 0.137 (0.034) | 0.265 (0.069) |  |
| JA-Ile | Herbivory | 0.421 (0.111) | 0.219 (0.048) | 0.545 (0.090) | 0.293 (0.081) | 0.474 (0.122) | 0.398 (0.122) |  |
|  | Drought | 0.184 (0.065) | 0.238 (0.053) | 0.179 (0.047) | 0.397 (0.077) | 0.128 (0.049) | 0.247 (0.040) |  |
|  | Combined | 0.646 (0.129) | 0.260 (0.053) | 0.639 (0.099) | 0.295 (0.075) | 0.528 (0.214) | 0.425 (0.113) |  |
| SA | Herbivory | 0.116 (0.052) | 0.244 (0.063) | 0.254 (0.083) | 0.418 (0.101) | 0.253 (0.127) | 0.284 (0.029) | Accession:<br>$\chi^2 = 19.59$;<br>$p = 0.02$ |
|  | Drought | 0.150 (0.090) | 0.265 (0.048) | 0.370 (0.074) | 0.354 (0.071) | 0.286 (0.108) | 0.402 (0.111) |  |
|  | Combined | 0.373 (0.111) | 0.246 (0.053) | 0.212 (0.096) | 0.462 (0.100) | 0.361 (0.102) | 0.381 (0.088) |  |
| ABA | Herbivory | 0.276 (0.063) | 0.193 (0.048) | 0.310 (0.081) | 0.301 (0.079) | 0.214 (0.054) | 0.294 (0.073) |  |
|  | Drought | 0.227 (0.101) | 0.193 (0.051) | 0.228 (0.077) | 0.295 (0.098) | 0.221 (0.084) | 0.356 (0.102) |  |
|  | Combined | 0.446 (0.110) | 0.239 (0.046) | 0.353 (0.098) | 0.356 (0.078) | 0.121 (0.011) | 0.334 (0.136) |  |
| IAA | Herbivory | 0.192 (0.070) | 0.199 (0.041) | 0.386 (0.089) | 0.251 (0.058) | 0.060 (0.032) | 0.188 (0.086) | Accession:<br>$\chi^2 = 14.05$;<br>$p = 0.02$ |
|  | Drought | 0.231 (0.048) | 0.427 (0.051) | 0.257 (0.112) | 0.462 (0.071) | 0.356 (0.087) | 0.284 (0.064) |  |
|  | Combined | 0.336 (0.112) | 0.411 (0.047) | 0.203 (0.154) | 0.336 (0.049) | 0.313 (0.066) | 0.336 (0.060) |  |
| Height | Herbivory | 0.187 (0.059) | 0.257 (0.063) | 0.210 (0.162) | 0.227 (0.050) | 0.089 (0.033) | 0.196 (0.069) |  |
|  | Drought | 0.096 (0.026) | 0.146 (0.030) | 0.257 (0.128) | 0.262 (0.057) | 0.147 (0.022) | 0.137 (0.020) |  |
|  | Combined | 0.119 (0.046) | 0.190 (0.036) | 0.373 (0.151) | 0.258 (0.049) | 0.141 (0.038) | 0.120 (0.063) |  |
| LN | Herbivory | 0.323 (0.079) | 0.130 (0.036) | 0.281 (0.063) | 0.338 (0.048) | 0.209 (0.080) | 0.126 (0.047) | Accession:<br>$\chi^2 = 27.37$;<br>$p < 0.001$ |
|  | Drought | 0.226 (0.057) | 0.172 (0.057) | 0.259 (0.079) | 0.404 (0.068) | 0.283 (0.050) | 0.088 (0.032) |  |
|  | Combined | 0.401 (0.076) | 0.230 (0.049) | 0.377 (0.150) | 0.413 (0.102) | 0.321 (0.077) | 0.198 (0.054) |  |
| LDM | Herbivory | 0.207 (0.103) | 0.091 (0.026) | 0.217 (0.056) | 0.246 (0.055) | 0.131 (0.045) | 0.172 (0.067) | Accession:<br>$\chi^2 = 17.41$;<br>$p = 0.004$ |
|  | Drought | 0.170 (0.049) | 0.156 (0.032) | 0.313 (0.035) | 0.343 (0.040) | 0.181 (0.079) | 0.244 (0.019) |  |
|  | Combined | 0.250 (0.062) | 0.164 (0.046) | 0.179 (0.058) | 0.242 (0.064) | 0.278 (0.078) | 0.188 (0.030) |  |
| SDM | Herbivory | 0.406 (0.106) | 0.484 (0.065) | 0.222 (0.053) | 0.362 (0.091) | 0.178 (0.073) | 0.199 (0.125) |  |
|  | Drought | 0.282 (0.040) | 0.204 (0.059) | 0.335 (0.123) | 0.287 (0.077) | 0.356 (0.134) | 0.307 (0.099) |  |
|  | Combined | 0.267 (0.062) | 0.321 (0.077) | 0.561 (0.089) | 0.436 (0.068) | 0.242 (0.114) | 0.412 (0.134) |  |
| ADM | Herbivory | 0.215 (0.104) | 0.160 (0.044) | 0.180 (0.057) | 0.274 (0.053) | 0.135 (0.048) | 0.187 (0.069) |  |
|  | Drought | 0.152 (0.045) | 0.127 (0.025) | 0.282 (0.072) | 0.289 (0.054) | 0.210 (0.066) | 0.239 (0.038) |  |
|  | Combined | 0.215 (0.043) | 0.191 (0.039) | 0.237 (0.047) | 0.266 (0.067) | 0.269 (0.083) | 0.173 (0.046) |  |
| RDM | Herbivory | 0.178 (0.040) | 0.173 (0.059) | 0.108 (0.065) | 0.180 (0.085) | 0.105 (0.086) | 0.178 (0.074) |  |
|  | Drought | 0.063 (0.011) | 0.106 (0.030) | 0.205 (0.039) | 0.194 (0.086) | 0.073 (0.017) | 0.141 (0.045) |  |
|  | Combined | 0.098 (0.034) | 0.079 (0.021) | 0.125 (0.046) | 0.260 (0.093) | 0.145 (0.075) | 0.215 (0.106) |  |
| TDM | Herbivory | 0.182 (0.045) | 0.159 (0.045) | 0.099 (0.049) | 0.132 (0.023) | 0.116 (0.050) | 0.155 (0.051) |  |
|  | Drought | 0.062 (0.018) | 0.095 (0.030) | 0.204 (0.047) | 0.166 (0.034) | 0.023 (0.010) | 0.108 (0.036) |  |
|  | Combined | 0.106 (0.036) | 0.099 (0.024) | 0.145 (0.036) | 0.210 (0.049) | 0.166 (0.091) | 0.196 (0.096) |  |
| RSR | Herbivory | 0.206 (0.064) | 0.168 (0.057) | 0.234 (0.096) | 0.276 (0.107) | 0.209 (0.072) | 0.219 (0.122) |  |
|  | Drought | 0.154 (0.042) | 0.112 (0.034) | 0.188 (0.075) | 0.281 (0.098) | 0.276 (0.069) | 0.296 (0.073) |  |
|  | Combined | 0.165 (0.048) | 0.117 (0.027) | 0.127 (0.067) | 0.320 (0.097) | 0.180 (0.044) | 0.162 (0.076) |  |
| LWC | Herbivory | 0.048 (0.024) | 0.034 (0.009) | 0.019 (0.005) | 0.035 (0.011) | 0.024 (0.012) | 0.060 (0.029) |  |
|  | Drought | 0.055 (0.016) | 0.061 (0.013) | 0.078 (0.021) | 0.064 (0.022) | 0.032 (0.014) | 0.063 (0.024) |  |
|  | Combined | 0.149 (0.037) | 0.083 (0.018) | 0.156 (0.047) | 0.086 (0.025) | 0.036 (0.012) | 0.078 (0.025) |  |

<sup>1</sup>RDPI: $(|x_c - x_s|)/(|x_c + x_s|)$ where $x_c$ and $x_s$ represent the different trait values of clones kept under control (c) and challenged (s) conditions
<sup>2</sup> Accessions represent propagations of each mother plant
<sup>3</sup> Full names are provided in Table 1

**Table 3.**
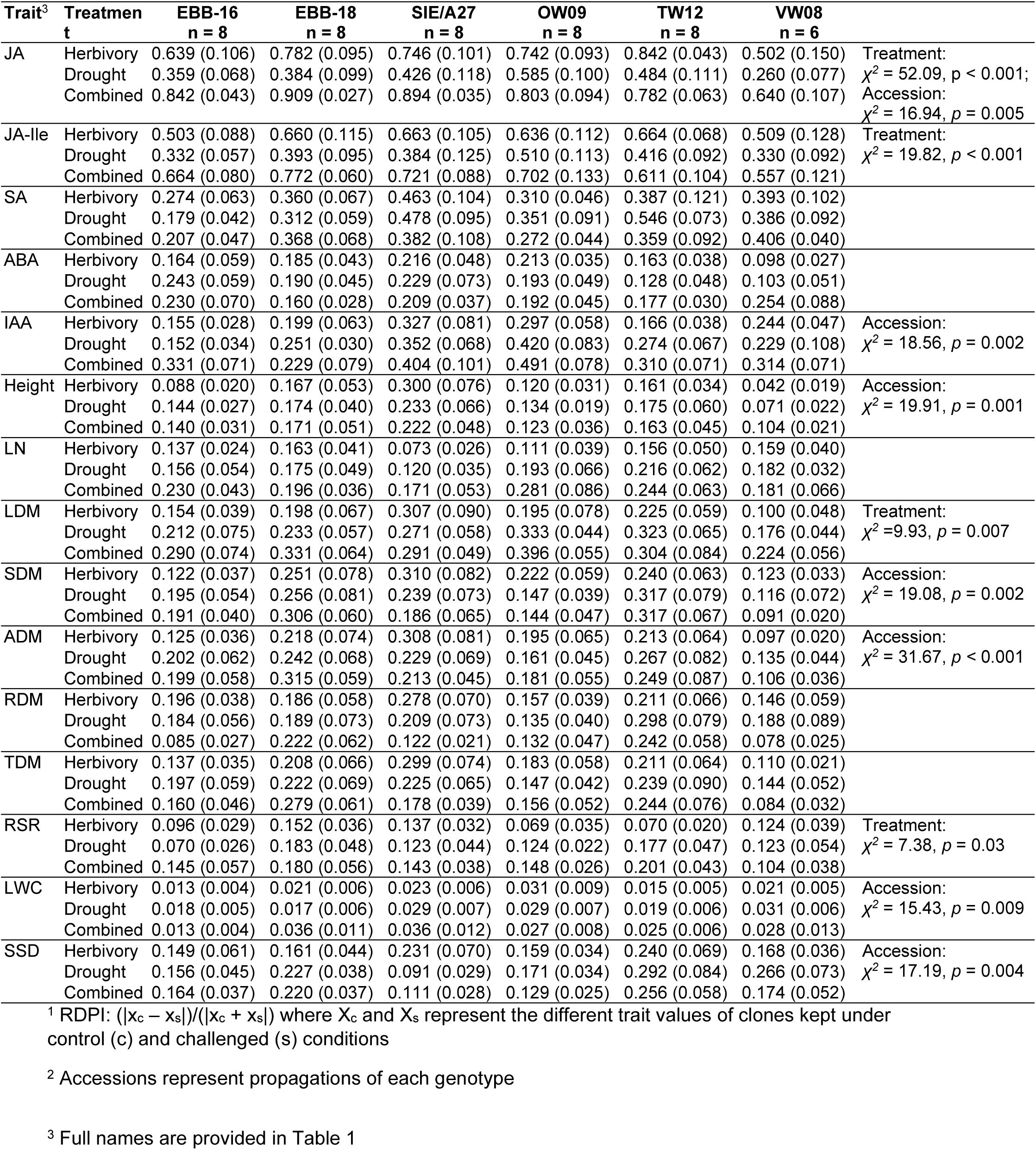
Means (standard deviations) of phenotypic plasticity (measured as RDPI^1^) of foliar phytohormone concentration and morphological traits of *Solanum dulcamara* of different accessions^2^, exposed to different environmental challenges (combined: drought and herbivory).

| Trait<sup>3</sup> | Treatment | EBB-16<br>n = 8 | EBB-18<br>n = 8 | SIE/A27<br>n = 8 | OW09<br>n = 8 | TW12<br>n = 8 | VW08<br>n = 6 |  |
| --- | --- | --- | --- | --- | --- | --- | --- | --- |
| JA | Herbivory | 0.639 (0.106) | 0.782 (0.095) | 0.746 (0.101) | 0.742 (0.093) | 0.842 (0.043) | 0.502 (0.150) | Treatment:<br>$\chi^2 = 52.09, p < 0.001$;<br>Accession:<br>$\chi^2 = 16.94, p = 0.005$ |
|  | Drought | 0.359 (0.068) | 0.384 (0.099) | 0.426 (0.118) | 0.585 (0.100) | 0.484 (0.111) | 0.260 (0.077) |  |
|  | Combined | 0.842 (0.043) | 0.909 (0.027) | 0.894 (0.035) | 0.803 (0.094) | 0.782 (0.063) | 0.640 (0.107) |  |
| JA-Ile | Herbivory | 0.503 (0.088) | 0.660 (0.115) | 0.663 (0.105) | 0.636 (0.112) | 0.664 (0.068) | 0.509 (0.128) | Treatment:<br>$\chi^2 = 19.82, p < 0.001$ |
|  | Drought | 0.332 (0.057) | 0.393 (0.095) | 0.384 (0.125) | 0.510 (0.113) | 0.416 (0.092) | 0.330 (0.092) |  |
|  | Combined | 0.664 (0.080) | 0.772 (0.060) | 0.721 (0.088) | 0.702 (0.133) | 0.611 (0.104) | 0.557 (0.121) |  |
| SA | Herbivory | 0.274 (0.063) | 0.360 (0.067) | 0.463 (0.104) | 0.310 (0.046) | 0.387 (0.121) | 0.393 (0.102) |  |
|  | Drought | 0.179 (0.042) | 0.312 (0.059) | 0.478 (0.095) | 0.351 (0.091) | 0.546 (0.073) | 0.386 (0.092) |  |
|  | Combined | 0.207 (0.047) | 0.368 (0.068) | 0.382 (0.108) | 0.272 (0.044) | 0.359 (0.092) | 0.406 (0.040) |  |
| ABA | Herbivory | 0.164 (0.059) | 0.185 (0.043) | 0.216 (0.048) | 0.213 (0.035) | 0.163 (0.038) | 0.098 (0.027) |  |
|  | Drought | 0.243 (0.059) | 0.190 (0.045) | 0.229 (0.073) | 0.193 (0.049) | 0.128 (0.048) | 0.103 (0.051) |  |
|  | Combined | 0.230 (0.070) | 0.160 (0.028) | 0.209 (0.037) | 0.192 (0.045) | 0.177 (0.030) | 0.254 (0.088) |  |
| IAA | Herbivory | 0.155 (0.028) | 0.199 (0.063) | 0.327 (0.081) | 0.297 (0.058) | 0.166 (0.038) | 0.244 (0.047) | Accession:<br>$\chi^2 = 18.56, p = 0.002$ |
|  | Drought | 0.152 (0.034) | 0.251 (0.030) | 0.352 (0.068) | 0.420 (0.083) | 0.274 (0.067) | 0.229 (0.108) |  |
|  | Combined | 0.331 (0.071) | 0.229 (0.079) | 0.404 (0.101) | 0.491 (0.078) | 0.310 (0.071) | 0.314 (0.071) |  |
| Height | Herbivory | 0.088 (0.020) | 0.167 (0.053) | 0.300 (0.076) | 0.120 (0.031) | 0.161 (0.034) | 0.042 (0.019) | Accession:<br>$\chi^2 = 19.91, p = 0.001$ |
|  | Drought | 0.144 (0.027) | 0.174 (0.040) | 0.233 (0.066) | 0.134 (0.019) | 0.175 (0.060) | 0.071 (0.022) |  |
|  | Combined | 0.140 (0.031) | 0.171 (0.051) | 0.222 (0.048) | 0.123 (0.036) | 0.163 (0.045) | 0.104 (0.021) |  |
| LN | Herbivory | 0.137 (0.024) | 0.163 (0.041) | 0.073 (0.026) | 0.111 (0.039) | 0.156 (0.050) | 0.159 (0.040) |  |
|  | Drought | 0.156 (0.054) | 0.175 (0.049) | 0.120 (0.035) | 0.193 (0.066) | 0.216 (0.062) | 0.182 (0.032) |  |
|  | Combined | 0.230 (0.043) | 0.196 (0.036) | 0.171 (0.053) | 0.281 (0.086) | 0.244 (0.063) | 0.181 (0.066) |  |
| LDM | Herbivory | 0.154 (0.039) | 0.198 (0.067) | 0.307 (0.090) | 0.195 (0.078) | 0.225 (0.059) | 0.100 (0.048) | Treatment:<br>$\chi^2 = 9.93, p = 0.007$ |
|  | Drought | 0.212 (0.075) | 0.233 (0.057) | 0.271 (0.058) | 0.333 (0.044) | 0.323 (0.065) | 0.176 (0.044) |  |
|  | Combined | 0.290 (0.074) | 0.331 (0.064) | 0.291 (0.049) | 0.396 (0.055) | 0.304 (0.084) | 0.224 (0.056) |  |
| SDM | Herbivory | 0.122 (0.037) | 0.251 (0.078) | 0.310 (0.082) | 0.222 (0.059) | 0.240 (0.063) | 0.123 (0.033) | Accession:<br>$\chi^2 = 19.08, p = 0.002$ |
|  | Drought | 0.195 (0.054) | 0.256 (0.081) | 0.239 (0.073) | 0.147 (0.039) | 0.317 (0.079) | 0.116 (0.072) |  |
|  | Combined | 0.191 (0.040) | 0.306 (0.060) | 0.186 (0.065) | 0.144 (0.047) | 0.317 (0.067) | 0.091 (0.020) |  |
| ADM | Herbivory | 0.125 (0.036) | 0.218 (0.074) | 0.308 (0.081) | 0.195 (0.065) | 0.213 (0.064) | 0.097 (0.020) | Accession:<br>$\chi^2 = 31.67, p < 0.001$ |
|  | Drought | 0.202 (0.062) | 0.242 (0.068) | 0.229 (0.069) | 0.161 (0.045) | 0.267 (0.082) | 0.135 (0.044) |  |
|  | Combined | 0.199 (0.058) | 0.315 (0.059) | 0.213 (0.045) | 0.181 (0.055) | 0.249 (0.087) | 0.106 (0.036) |  |
| RDM | Herbivory | 0.196 (0.038) | 0.186 (0.058) | 0.278 (0.070) | 0.157 (0.039) | 0.211 (0.066) | 0.146 (0.059) |  |
|  | Drought | 0.184 (0.056) | 0.189 (0.073) | 0.209 (0.073) | 0.135 (0.040) | 0.298 (0.079) | 0.188 (0.089) |  |
|  | Combined | 0.085 (0.027) | 0.222 (0.062) | 0.122 (0.021) | 0.132 (0.047) | 0.242 (0.058) | 0.078 (0.025) |  |
| TDM | Herbivory | 0.137 (0.035) | 0.208 (0.066) | 0.299 (0.074) | 0.183 (0.058) | 0.211 (0.064) | 0.110 (0.021) |  |
|  | Drought | 0.197 (0.059) | 0.222 (0.069) | 0.225 (0.065) | 0.147 (0.042) | 0.239 (0.090) | 0.144 (0.052) |  |
|  | Combined | 0.160 (0.046) | 0.279 (0.061) | 0.178 (0.039) | 0.156 (0.052) | 0.244 (0.076) | 0.084 (0.032) |  |
| RSR | Herbivory | 0.096 (0.029) | 0.152 (0.036) | 0.137 (0.032) | 0.069 (0.035) | 0.070 (0.020) | 0.124 (0.039) | Treatment:<br>$\chi^2 = 7.38, p = 0.03$ |
|  | Drought | 0.070 (0.026) | 0.183 (0.048) | 0.123 (0.044) | 0.124 (0.022) | 0.177 (0.047) | 0.123 (0.054) |  |
|  | Combined | 0.145 (0.057) | 0.180 (0.056) | 0.143 (0.038) | 0.148 (0.026) | 0.201 (0.043) | 0.104 (0.038) |  |
| LWC | Herbivory | 0.013 (0.004) | 0.021 (0.006) | 0.023 (0.006) | 0.031 (0.009) | 0.015 (0.005) | 0.021 (0.005) | Accession:<br>$\chi^2 = 15.43, p = 0.009$ |
|  | Drought | 0.018 (0.005) | 0.017 (0.006) | 0.029 (0.007) | 0.029 (0.007) | 0.019 (0.006) | 0.031 (0.006) |  |
|  | Combined | 0.013 (0.004) | 0.036 (0.011) | 0.036 (0.012) | 0.027 (0.008) | 0.025 (0.006) | 0.028 (0.013) |  |
| SSD | Herbivory | 0.149 (0.061) | 0.161 (0.044) | 0.231 (0.070) | 0.159 (0.034) | 0.240 (0.069) | 0.168 (0.036) | Accession:<br>$\chi^2 = 17.19, p = 0.004$ |
|  | Drought | 0.156 (0.045) | 0.227 (0.038) | 0.091 (0.029) | 0.171 (0.034) | 0.292 (0.084) | 0.266 (0.073) |  |
|  | Combined | 0.164 (0.037) | 0.220 (0.037) | 0.111 (0.028) | 0.129 (0.025) | 0.256 (0.058) | 0.174 (0.052) |  |

<sup>1</sup> RDPI: $(|x_c - x_s|)/(|x_c + x_s|)$ where $x_c$ and $x_s$ represent the different trait values of clones kept under control (c) and challenged (s) conditions
<sup>2</sup> Accessions represent propagations of each genotype
<sup>3</sup> Full names are provided in Table 1

**Table 4.** Means (standard deviations) of phenotypic plasticity (measured as RDPI^1^) of foliar phytohormone concentration and morphological traits of *Populus nigra* of different accessions^2^, exposed to different environmental challenges (combined: drought and herbivory).^1^RDPI: (|x_c_ – x_s_|)/(|x_c_ + x_s_|) where X_c_ and X_s_ represent the different trait values of clones kept under control (c) and challenged (s) conditions.

| Trait<sup>3</sup> | Treatment | BW_R14<br>n = 4 | BW_R19<br>n = 4 | BW_R20<br>n = 4 | BW_R25<br>n = 4 | BW_R30<br>n = 4 | BW_R32<br>n = 4 |  |
| --- | --- | --- | --- | --- | --- | --- | --- | --- |
| JA | Herbivory | 0.604 (0.151) | 0.722 (0.074) | 0.643 (0.156) | 0.605 (0.122) | 0.818 (0.055) | 0.691 (0.072) | Treatment:<br>$X^2 = 25.88$,<br>$p < 0.001$ |
|  | Drought | 0.296 (0.087) | 0.255 (0.028) | 0.418 (0.128) | 0.120 (0.055) | 0.389 (0.050) | 0.392 (0.148) |  |
|  | Combined | 0.307 (0.165) | 0.574 (0.171) | 0.555 (0.067) | 0.583 (0.133) | 0.481 (0.114) | 0.467 (0.157) |  |
| JA-Ile | Herbivory | 0.514 (0.201) | 0.553 (0.161) | 0.393 (0.206) | 0.520 (0.134) | 0.746 (0.079) | 0.409 (0.135) |  |
|  | Drought | 0.376 (0.164) | 0.388 (0.211) | 0.351 (0.102) | 0.360 (0.117) | 0.612 (0.140) | 0.566 (0.152) |  |
|  | Combined | 0.357 (0.150) | 0.420 (0.147) | 0.232 (0.022) | 0.622 (0.140) | 0.451 (0.078) | 0.405 (0.081) |  |
| SA | Herbivory | 0.330 (0.085) | 0.237 (0.093) | 0.231 (0.055) | 0.193 (0.029) | 0.130 (0.100) | 0.160 (0.042) |  |
|  | Drought | 0.412 (0.153) | 0.328 (0.113) | 0.360 (0.060) | 0.193 (0.100) | 0.217 (0.062) | 0.170 (0.024) |  |
|  | Combined | 0.228 (0.106) | 0.302 (0.081) | 0.287 (0.088) | 0.317 (0.153) | 0.309 (0.091) | 0.135 (0.051) |  |
| ABA | Herbivory | 0.516 (0.138) | 0.342 (0.063) | 0.359 (0.083) | 0.523 (0.181) | 0.218 (0.074) | 0.493 (0.088) | Treatment:<br>$X^2 = 61.01$,<br>$p < 0.001$ |
|  | Drought | 0.922 (0.024) | 0.747 (0.158) | 0.898 (0.036) | 0.952 (0.010) | 0.794 (0.069) | 0.646 (0.125) |  |
|  | Combined | 0.916 (0.042) | 0.822 (0.068) | 0.718 (0.180) | 0.938 (0.018) | 0.733 (0.083) | 0.797 (0.072) |  |
| IAA | Herbivory | 0.108 (0.040) | 0.116 (0.039) | 0.343 (0.086) | 0.233 (0.057) | 0.229 (0.056) | 0.109 (0.043) | Accession:<br>$X^2 = 56.89$,<br>$p < 0.001$ |
|  | Drought | 0.145 (0.022) | 0.223 (0.048) | 0.459 (0.076) | 0.067 (0.028) | 0.211 (0.065) | 0.170 (0.040) |  |
|  | Combined | 0.157 (0.062) | 0.146 (0.055) | 0.486 (0.084) | 0.131 (0.039) | 0.244 (0.065) | 0.124 (0.049) |  |
| Height | Herbivory | 0.077 (0.027) | 0.017 (0.007) | 0.047 (0.025) | 0.075 (0.025) | 0.043 (0.018) | 0.051 (0.014) |  |
|  | Drought | 0.068 (0.013) | 0.027 (0.013) | 0.074 (0.036) | 0.095 (0.046) | 0.072 (0.021) | 0.111 (0.014) |  |
|  | Combined | 0.067 (0.019) | 0.063 (0.012) | 0.072 (0.028) | 0.077 (0.041) | 0.058 (0.024) | 0.059 (0.005) |  |
| LN | Herbivory | 0.033 (0.017) | 0.103 (0.047) | 0.035 (0.016) | 0.052 (0.014) | 0.055 (0.023) | 0.093 (0.020) |  |
|  | Drought | 0.041 (0.018) | 0.141 (0.065) | 0.074 (0.025) | 0.076 (0.043) | 0.033 (0.017) | 0.082 (0.017) |  |
|  | Combined | 0.035 (0.021) | 0.157 (0.079) | 0.091 (0.032) | 0.154 (0.036) | 0.076 (0.024) | 0.077 (0.023) |  |
| LDM | Herbivory | 0.141 (0.051) | 0.079 (0.038) | 0.052 (0.023) | 0.133 (0.055) | 0.030 (0.009) | 0.044 (0.022) |  |
|  | Drought | 0.093 (0.023) | 0.175 (0.051) | 0.116 (0.056) | 0.153 (0.059) | 0.098 (0.041) | 0.058 (0.028) |  |
|  | Combined | 0.122 (0.035) | 0.106 (0.035) | 0.137 (0.031) | 0.085 (0.055) | 0.093 (0.029) | 0.090 (0.020) |  |
| SDM | Herbivory | 0.234 (0.023) | 0.081 (0.031) | 0.094 (0.051) | 0.166 (0.043) | 0.037 (0.010) | 0.070 (0.029) |  |
|  | Drought | 0.106 (0.027) | 0.080 (0.023) | 0.089 (0.039) | 0.207 (0.082) | 0.107 (0.069) | 0.094 (0.019) |  |
|  | Combined | 0.136 (0.013) | 0.113 (0.026) | 0.091 (0.016) | 0.132 (0.063) | 0.139 (0.029) | 0.063 (0.038) |  |
| ADM | Herbivory | 0.167 (0.040) | 0.080 (0.029) | 0.066 (0.017) | 0.146 (0.032) | 0.017 (0.009) | 0.041 (0.028) | Accession:<br>$X^2 = 22.47$,<br>$p < 0.001$ |
|  | Drought | 0.086 (0.025) | 0.095 (0.033) | 0.108 (0.047) | 0.148 (0.076) | 0.094 (0.055) | 0.050 (0.024) |  |
|  | Combined | 0.126 (0.020) | 0.081 (0.035) | 0.079 (0.025) | 0.106 (0.054) | 0.035 (0.011) | 0.042 (0.015) |  |
| RDM | Herbivory | 0.076 (0.011) | 0.129 (0.025) | 0.106 (0.056) | 0.148 (0.057) | 0.021 (0.005) | 0.125 (0.017) |  |
|  | Drought | 0.080 (0.028) | 0.137 (0.059) | 0.139 (0.030) | 0.177 (0.087) | 0.112 (0.054) | 0.161 (0.050) |  |
|  | Combined | 0.083 (0.037) | 0.130 (0.048) | 0.066 (0.027) | 0.157 (0.056) | 0.194 (0.055) | 0.128 (0.011) |  |
| TDM | Herbivory | 0.134 (0.028) | 0.065 (0.033) | 0.077 (0.026) | 0.146 (0.040) | 0.018 (0.005) | 0.067 (0.021) |  |
|  | Drought | 0.070 (0.022) | 0.068 (0.027) | 0.117 (0.038) | 0.151 (0.084) | 0.094 (0.056) | 0.093 (0.029) |  |
|  | Combined | 0.106 (0.028) | 0.078 (0.032) | 0.046 (0.018) | 0.125 (0.052) | 0.085 (0.029) | 0.050 (0.019) |  |
| RSR | Herbivory | 0.092 (0.038) | 0.108 (0.052) | 0.071 (0.035) | 0.063 (0.010) | 0.018 (0.008) | 0.095 (0.036) |  |
|  | Drought | 0.092 (0.035) | 0.139 (0.094) | 0.080 (0.021) | 0.045 (0.023) | 0.056 (0.008) | 0.119 (0.038) |  |
|  | Combined | 0.061 (0.019) | 0.121 (0.058) | 0.128 (0.041) | 0.060 (0.031) | 0.191 (0.057) | 0.133 (0.027) |  |
| LWC | Herbivory | 0.021 (0.008) | 0.014 (0.006) | 0.012 (0.005) | 0.058 (0.027) | 0.016 (0.004) | 0.020 (0.006) |  |
|  | Drought | 0.014 (0.006) | 0.039 (0.014) | 0.026 (0.010) | 0.017 (0.008) | 0.037 (0.007) | 0.033 (0.009) |  |
|  | Combined | 0.026 (0.010) | 0.034 (0.009) | 0.048 (0.020) | 0.082 (0.053) | 0.047 (0.014) | 0.047 (0.019) |  |
| SSD | Herbivory | 0.164 (0.060) | 0.138 (0.053) | 0.193 (0.038) | 0.163 (0.058) | 0.267 (0.124) | 0.219 (0.057) |  |
|  | Drought | 0.158 (0.035) | 0.178 (0.046) | 0.198 (0.093) | 0.246 (0.106) | 0.189 (0.055) | 0.193 (0.060) |  |
|  | Combined | 0.157 (0.072) | 0.066 (0.031) | 0.249 (0.094) | 0.072 (0.036) | 0.142 (0.062) | 0.111 (0.051) |  |

<sup>2</sup>Accessions represent propagations of each genotype
<sup>3</sup> Full names are provided in Table 1

### Plant morphological traits mostly respond to treatments in a species-specific manner

Similar to foliar phytohormones, morphological traits differed among species (Table 1); therefore, they also contributed to the grouping of species, with *T. vulgare* tending to exhibit a higher root dry mass, total dry mass, and root-shoot ratio compared with *S. dulcamara*, whereas *S. dulcarama* was characterized by a higher shoot height, leaf number, leaf water content, stem dry mass, and aboveground dry mass. Samples of *P. nigra* were distinguished primarily by a higher leaf dry mass (Fig. 2A).

In general, shoot height, leaf number, leaf dry mass, leaf water content, and root-shoot ratio were significantly impacted by treatment in all three species. Meanwhile, root dry mass remained unchanged among treatments in all three species (Table 1, Fig. S5J-L). Moreover, several species-specific patterns were found. Within *T. vulgare*, morphological traits also contributed to the separation of the control and combined treatment (Fig. 2B). In particular, aboveground dry mass was reduced under drought and the combined treatment in *T. vulgare* (Fig. 3G), which was mirrored in an increased root-shoot ratio (Fig. 3J). Similarly, in *S. dulcamara*, the root-shoot ratio was positively associated with the combined treatment, while leaf number, leaf dry mass, shoot height, aboveground dry mass, and total dry mass were reduced by the combined treatment (Fig. 2C). The aboveground dry mass was also lower under the drought and the combined treatment than in the controls in *S. dulcamara* (Fig. 3H). In contrast to *T. vulgare*, the root-shoot ratio was only enhanced under the combined treatment in *S. dulcamara* (Fig. 3K). In contrast, *P. nigra* exhibited a higher root–shoot ratio and specific stem density under the drought and the combined treatment, whereas shoot height, leaf number, and leaf water content were higher under the herbivory and the control treatment (Fig. 2D). Only in *P. nigra*, aboveground dry mass remained unchanged among treatments (Fig 3I).

Apart from treatment effects, across all three species all morphological traits were significantly affected by accession, except leaf water content in *T. vulgare* and specific stem density in *P. nigra* (Table 1). However, there were no effects of sex on any morphological trait of *P. nigra* (Table 1).

Patterns of phenotypic plasticity (RDPI) for morphological traits varied among species. In *T. vulgare*, accession affected the RDPIs of leaf number and leaf dry mass (Table 2). In *S. dulcamara*, treatment significantly influenced the RDPI of the leaf dry mass and the root–shoot ratio, with higher RDPI values under the combined treatment than under single challenges, except in two accessions (SIE/A27 and TW12) for leaf dry mass, and one (VW08) for root-shoot ratio. Beyond the treatment effect, the RDPIs of morphological traits in *S. dulcamara* were more pronouncedly shaped by accession, affecting shoot height, stem dry mass, aboveground dry mass, leaf water content, and specific stem density (Table 3). In *P. nigra*, none of the RDPIs of the morphological traits was affected by treatment, while only the aboveground dry mass was affected by accession, and the leaf water content was impacted by sex (Tables 4, S7).

### Drought drecreased herbivore performance in *P. nigra* only

The performance of the herbivores used in the herbivory and combined treatment was assessed by measuring their relative growth rate. Growth rates were comparable on plants of both treatments for larvae of *Spodoptera exigua* on both *T. vulgare* (*F* = 2.09, *p* = 0.15, Fig. 4A) and *S. dulcamara*, although there was a slight trend for a reduction under the combined treatment in *S. dulcamara* (*F* = 3.46, *p* = 0.07, Fig. 4B). On *P. nigra* plants, average herbivore growth rates were significantly lower for larvae of *Lymantria dispar* fed on drought-challenged plants (*F* = 16.54*, p* < 0.001, Fig. 4C). There was no difference in leaf area loss between plants of the herbivory and combined treatment in both *S. dulcamara* and *P. nigra* (Fig. S7).

**Fig. 4.**
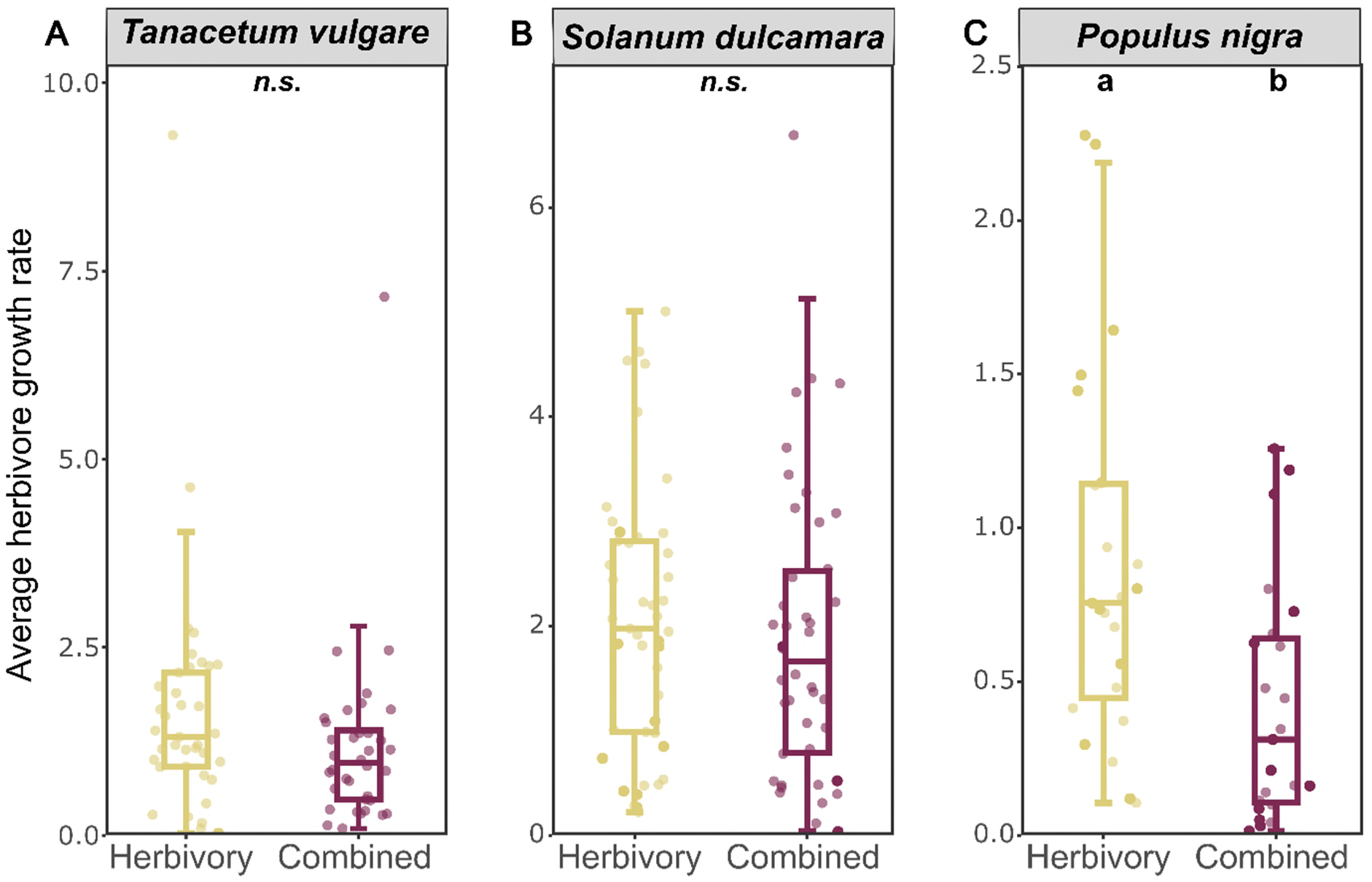
Average herbivore growth rate of *Spodoptera exigua* on *Tanacetum vulgare* (A), *Solanum dulcamara* (B); and of *Lymanthria dispar* on *Populus nigra* (C), respectively, during exposure time to plants subjected to herbivory and the combined treatment (herbivory and drought). Data are presented as boxplots, with medians, interquartile ranges (IQR, boxes), and whiskers extending to the most extreme values with max. 1.5 times the IQR. Individual values are plotted as dots; *n* = 48 per treatment for *T. vulgare* and *S. dulcamara*, n = 24 per treatment for *P. nigra*. Different letters indicate statistically significant differences (Mann-Whitney U -test, *p* < 0.05); *n.s.*: not significant.

### Phytohormones potentially mediate treatment-effects on plant morphological traits

Most of the final SEMs for *T. vulgare*, *S. dulcamara*, and *P. nigra* provided appropriate representations of the overall correlation structure of the measured variables, as indicated by non-significant goodness-of-fit tests, except for treatment and sex on foliar phytohormone and morphological traits in *P. nigra* (Table S8). In all three species, the effect of treatment on root dry mass was consistently mediated by ABA concentrations, although different relationships were observed among species.

In *T. vulgare*, JA-Ile, ABA, and IAA concentrations responded directly to treatment, and all phytohormones were affected by accession (Fig. 5A). ABA positively affected (enhanced) root dry mass, whereas IAA negatively influenced (reduced) root dry mass. In addition, JA negatively affected root dry mass but positively influenced leaf number. SA negative affected stem dry mass. Treatment as well as accession also directly affected leaf number, leaf dry mass, and plant height, while treatment additionally affected stem dry mass and leaf water content, and accession additionally affected root dry mass (Fig. 5B). Beyond these direct effects, JA emerged as a key mediator influencing leaf number, root dry mass, and root-shoot ratio (Fig. 5C).

**Fig. 5.**
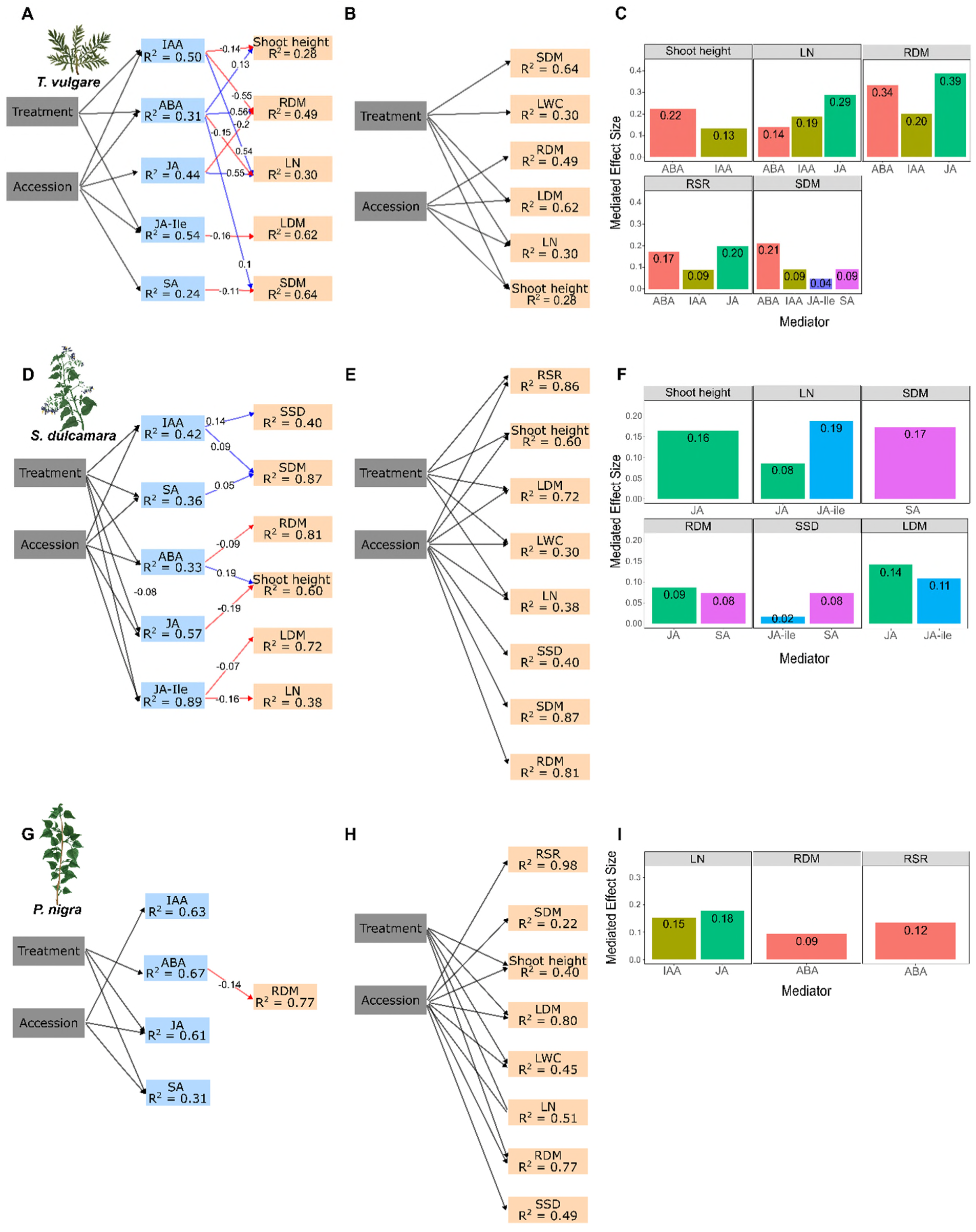
Structural equation models (SEMs) showing how foliar phytohormones may influence morphological responses to treatments in *Tanacetum vulgare*, *Solanum dulcamara*, and *Populus nigra*. Significant and direct relationships between treatment and accession on foliar phytohormone concentrations and between phytohormones and morphological traits in *T. vulgare* (A), *S. dulcamara* (D), and *Populus nigra* (G). Numbers on each arrow represent standardized effect sizes. Blue arrows indicate positive relationships, red arrows indicate negative ones. Direct effects of treatment and accession on morphological traits in *T. vulgare* (B), *S. dulcamara* (E), *P. nigra* (H). All SEMs fitted data well (*T. vulgare*: Fisher’s C = 59.61, *df* = 46, *p* = 0.09; *S. dulcamara*: Fisher’s C = 70.76, *df* = 56, *p* = 0.09; *P. nigra*: Fisher’s C = 80.005, *df* = 86, *p* = 0.66). Coefficients of determination (R²) for each endogenous variable are shown in the corresponding boxes. For each SEM, the magnitudes of phytohormone mediator effects (i.e., all indirect pathways linking treatment or accession to morphological traits) are shown as bar plots (C: *T. vulgare*; F: *S. dulcamara*; I: *P. nigra*). Bootstrapped mediator effects are presented as absolute values to allow comparison of effect strength regardless of direction. The morphological traits are: LN: leaf number; LDM: leaf dry mass; SDM: stem dry mass; RDM: root dry mass; RSR: root-shoot ratio; LWC: leaf water content; SSD: specific-stem-density.

In *S. dulcamara*, all phytohormones were directly affected by both treatment and accession (Fig. 5D). Among the direct relationships between phytohormones and morphological traits, positive effects were observed between SA and stem dry mass, ABA and shoot height, and IAA with both stem dry mass and specific stem density. In contrast, JA showed negative effects on shoot height, and JA-Ile on leaf number and leaf dry mass, while ABA exerted a negative effect on root dry mass. Treatment directly influenced shoot height, leaf number, leaf dry mass, and root–shoot ratio, whereas accessions differently affected all morphological traits included in the SEM (Fig.5E). JA and/or JA-Ile acted as key mediators of variation in shoot height, leaf number, root dry mass, specific stem density, and leaf dry mass. In addition, SA influenced root dry mass, specific stem density, and stem dry mass (Fig. 5F).

In *P. nigra*, the SEM including treatment and accession provided a better fit to the data than the SEM including treatment and sex (accession: AICc = 2464.29; sex: AICc = 2649.72; Figs. 5G and S8). Treatment exerted direct effects on ABA, JA, and SA concentrations, whereas accession directly affected IAA, JA, and SA. Regarding direct relationships between phytohormones and morphological traits, only ABA negatively impacted with root dry mass (Fig. 5G). Treatment and accession also directly influenced most morphological traits, with root-shoot ratio, stem dry mass, and specific stem density being affected only by accession (Fig. 5H). Beyond these direct effects, JA and IAA mediated variation in leaf number, while ABA influenced root dry mass and root–shoot ratio (Fig. 5I).

## Discussion

Plants are known to respond to the increasingly common environmental challenges, such as drought and herbivory, by adjusting their phytohormone levels, morphological traits, and metabolic profiles [1, 19]. However, studies applying both challenges individually and in combination and testing responses of plants of different families and with different growth forms under comparable experimental conditions, thus allowing for direct comparisons, remain scarce. To address this gap, we investigated responses of phytohormones, morphological traits, and their phenotypic plasticity to comparable environmental challenges in *T. vulgare*, *S. dulcamara*, and *P. nigra*, representing an herbaceous, a woody vine, and a tree species, respectively. We found both shared and species-specific responses to the different treatments, with the tree species showing particularly distinct response patterns compared to the other two species. Notably, many traits were also influenced within species by accession, indicating that environmental conditions and genetic background jointly shape plant traits and their plasticity across growth forms. Moreover, species-specific patterns were revealed in the regulation of morphological traits by phytohormones under both single and combined challenges.

### General and species-specific responses in foliar phytohormones

Plants exposed to drought alone or in combination with herbivory consistently exhibited elevated ABA concentrations across all three species as expected, whereas IAA increased most consistently under the combined treatment. ABA is widely regarded as a primary regulator of plant responses to drought, whereas IAA can contribute to growth adjustment and water-balance regulation depending on species and context [9]. ABA was also induced by herbivory alone in *S. dulcamara* and *P. nigra*, but not in *T. vulgare*. Herbivory can decrease leaf water potential [49], which leads to an induction of ABA, as shown in several species [16, 21].

In response to herbivory, we expected an activation of the jasmonate signaling cascade, including JA and its conjugate JA-Ile, as is commonly reported [4, 50]. Indeed, JA and JA-Ile were significantly induced due to herbivory in *S. dulcamara* and *P. nigra* in line with our expectation. However, in *T. vulgare,* JA remained unchanged. This unexpected pattern may reflect comparatively high constitutive JA concentrations in *T. vulgare*, which may have limited a further measurable increase, or the herbivore damage was insufficient to alter JA concentration. Alternatively, herbivory in *T. vulgare* may have primarily promoted the conversion of JA into its bioactive conjugate JA-Ile, as is indicated by the pronounced induction of JA-Ile under both the herbivory and the combined treatment in this species. In addition, the potential antagonistic interactions between signaling of ABA and JA may have contributed to the unchanged JA concentrations observed in *T. vulgare* [4]. SA also showed species-specific treatment responses, with a significant overall treatment effect in *S. dulcamara* but no clear pairwise differences, and a distinct response in *P. nigra*. In *P. nigra*, SA concentrations were higher in plants of the combined treatment compared to the control. Beyond its primary role in plant immunity, SA also regulates plant responses to abiotic challenges depending on species [51]. Environmental challenges also affected SA in *P. tomentosa*, likely reflecting SA-mediated systemic acquired resistance, which is characteristic for slow-growing trees [52].

### Species-specific impacts on morphological plant traits, with comparatively weak biomass responses in *P. nigra*

Aboveground dry mass was significantly reduced by the two-week drought treatment and especially by the combined challenge in the two fast-growing species, *T. vulgare* and *S. dulcamara*, in line with our expectations. Drought is widely known to constrain plant growth by limiting carbon assimilation through stomatal closure and by directly impairing photosynthetic processes, often resulting in reduced biomass accumulation [53]. In contrast to the other two species, total aboveground and stem dry mass remained unaffected by treatment in *P. nigra*, but shoot height and leaf number were nevertheless significantly reduced under drought and the combined treatment. The absence of a drought effect on stem dry mass is consistent with previous observations in *P. nigra* following three weeks of drought exposure [52]. However, a reduced stem dry mass was found in other *Populus* species under longer or more severe drought conditions [54]. These plants were also much higher (> 1 m) than our poplar seedlings (∼ 0.4 m). This suggests that a longer duration or higher intensity of drought may be required to elicit changes in stem dry mass in *P. nigra* and/or plants may respond stronger at a more mature developmental stage. In the present study, the lack of change in stem dry mass of *P. nigra* may also reflect compensatory structural adjustment, because specific stem density increased under drought and the combined challenge, although shoot height declined. A higher stem density has been linked to higher drought tolerance across woody plant species, suggesting a structural adjustment that enhances resistance to water stress [55]. Woody and relatively slow-growing species often exhibit greater resistance to climatic extremes through more conservative growth strategies compared to fast-growing species [56]. In addition, they grow taller, live longer and show overall more resource investment in supportive structures, such as stems [57, 58].

Herbivory alone did not significantly affect aboveground dry mass in any of the three species in line with other studies [59, 60]. This may indicate that the imposed feeding pressure exerted by three larvae per plant was relatively low, or that plants were able to compensate for tissue loss through regrowth. Such compensatory growth responses are commonly observed as long as water and nutrient availability are still sufficient [61].

Belowground, drought is generally expected to promote higher investment in roots relative to shoots in order to optimize water uptake under limiting conditions [53]. Contrary to this expectation, root dry mass did not increase under drought or the combined challenge in any of the focal species. One possible explanation is the relatively small pot volume (2.2 L) used in our experiment, which may have constrained root growth and led to early saturation of belowground space. Nevertheless, the root-shoot ratio increased across the fast-growing species *T. vulgare* and *S. dulcamara* under these treatments, indicating a shift in biomass allocation driven primarily by reduced aboveground growth rather than enhanced root production. The pattern was particularly evident in *S. dulcamara*, where root dry mass remained low and was therefore less likely to have been limited by pot capacity. This pattern suggests that aboveground organs are more sensitive to short-term water limitation than belowground tissues, consistent with previous findings across plants of different functional types [62].

Despite significant treatment effects on mean trait values, changes in phenotypic plasticity were less pronounced. Because RDPI quantifies relative differences in trait expression between challenge and control environments, it reflects responsiveness rather than absolute shifts in trait means [48]. Consequently, significant shifts in trait means do not necessarily result in increased RDPI values. Nonetheless, species-specific patterns in plasticity of morphological traits were evident, with RDPI values of two traits being affected by treatment in *S. dulcamara*, one in *T. vulgare* and none in *P. nigra*, indicating a more conservative phenotypic response to environmental challenges in the latter.

### Intra-specific variation of responses to challenges in foliar phytohormones and morphological traits

Accessions correspond to distinct genotypes in *S. dulcamara* and *P. nigra*, and likely also represent differences in genetic background in *T. vulgare*. Responses in phytohormones and morphological traits differed among accessions across all focal species to different degrees. All measured traits were affected by accession in *S. dulcamara,* and nearly all traits except leaf water content varied among accessions in *T. vulgare*, indicating substantial intraspecific variation in both species. In fact, both species are known to occur across a broad ecological range and to exhibit pronounced plasticity [63, 64]. Only two phytohormones, but almost all morphological traits, were affected by accession in *P. nigra*. Substantial genotypic variation in the responsiveness of morphological and biochemical traits to drought and herbivory has also been documented in *Solanum tuberosum* [65], a congeneric species of *S. dulcamara*, as well as in members of the Salicaceae [66], including *P. nigra* [67]. Moreover, male trees of *P. nigra* exhibited higher responsiveness to both ABA and SA under challenging conditions than female trees, reflecting sex-specific differences in hormone signaling. Higher ABA induction was likewise found in male compared to female *Salix rehderiana* [68], suggesting that males may prioritize rapid hormonal adjustments to regulate water balance and defense, whereas females may maintain a more stable state.

### Species-specific impacts of phytohormones on morphological traits

Based on the SEMs, ABA acted as key intermediate variable linking treatment to variation in root dry mass in all three species, consistent with its central role in coordinating plant responses to environmental challenges [9, 69]. However, the nature of the interaction between ABA and IAA differed markedly between species. The auxin IAA exhibited an antagonistic interaction with ABA in *T. vulgare* on root dry mass, whereas no interaction was observed in *S. dulcamara*, highlighting that auxin–ABA crosstalk is species-specific [70]. Interestingly, SEM supported ABA promoted root growth in *T. vulgare*, but not in *S. dulcamara* and *P. nigra* where ABA was negatively correlated with root dry mass. This contrasting pattern may be partly explained by the higher baseline ABA concentrations observed in *T. vulgare* when compared with the other two species. Although ABA is often regarded as a growth inhibitor, its effects are concentration-dependent, with low concentrations stimulating root growth and higher concentrations repressing it [71]. We are aware that we only measured end points of phytohormone concentrations, so we cannot depict trajectories. Also, manipulations of phytohormone concentrations may be needed to test their direct effects. Nevertheless, SEMs provide a useful framework for identifying biologically plausible pathways, although causal interpretation requires caution [72].

The importance of JA- and JA-Ile-mediated effects on morphological traits also varied markedly among *T. vulgare*, *S. dulcamara*, and *P. nigra*. Finally, SA, as the defensive phytohormone primarily involved in response to phloem sap feeders and biotrophic pathogens [73], differed among accessions only in *S. dulcamara* and *P. nigra*. These findings highlight the species-specificity in phytohormonal crosstalk, resulting in divergent growth outcomes among plants [74]. They also indicate that genetic variation shapes plant phenotype partly through phytohormone-mediated pathways [75].

### Ecological implications

By applying comparable drought and herbivory treatments across multiple plant species, this study provides a complementary, experimental perspective to existing meta-analyses that synthesize plant responses across diverse systems [76, 77]. Meta-analyses undeniably provide an efficient means for synthesizing enormous ecological data and enable comparisons of effect sizes, such as those associated with drought, herbivory, and their combination. However, the robustness and reliability of meta-analytic outcomes depend critically on the use of standardized metrics and well-defined experimental conditions [78]. In this context, the absence of a clear and widely accepted definition of drought challenge or stress remains a major limitation [79]. Studying plant responses under comparable experimental conditions allowed us to reveal general, species-specific, and intraspecific accession-patterns in phytohormonal and morphological responses to single and combined environmental challenges. Beyond phytohormones and morphological traits, future research should extend the current framework by integrating transcriptomic and metabolomic approaches to uncover the molecular mechanisms underlying these phenotypic and hormonal patterns. Comparative transcriptome analyses could reveal how stress signaling networks and phytohormone-related genes are differentially regulated across species under identical challenges, while metabolite profiling could capture downstream biochemical adjustments and resource reallocation.

## Conclusion

This study applied comparable abiotic and biotic challenges, namely drought and herbivory, alone and in combination to three plants species to evaluate plant phytohormonal and morphological responses, as well as intraspecific variation in responses. Our results demonstrate some common responses across species, but also species-specific responses, especially in the magnitude of phytohormonal changes. In addition to treatment-specific effects, accession-specific responses were found, indicating variation in phenotypic plasticity within species. With ever increasing environmental change, the complexity of plant responses must be considered at multiple levels.

## List of abbreviations

JA: jasmonic acid
JA-Ile: jasmonoyl-isoleucine
SA: salicylic acid
ABA: abscisic acid
IAA: indole acetate acid
ET: ethylene
LN: leaf number
LDM: leaf dry mass
SDM: stem dry mass
ADM: aboveground dry mass
RDM: root dry mass
TDM: total dry mass
RSR: root-shoot ratio
LWC: leaf water content
SSD: specific stem density
RDPI: relative distance plasticity index
RDA: redundancy analysis
LM: linear model
glmmTMB: general linear mixed model template model builder
ANOVA: analysis of variance
SEM: structural equation model;

## Declaration

### Ethics approval and consent to participate

Not applicable.

### Consent for publication

Not applicable.

### Availability of data and materials

The datasets used and/or analyzed during the current study are available from the corresponding author and will be made available via DataPlant upon publication of this manuscript.

### Authors’ contributions

AS, AB, TD, RH, RRJ, SBU, NMvD, WWW, JPS, and CM conceived the ideas and proposed the conception of the experimental design. XX and JBW contributed to the experimental design, coordinated and conducted the climate chamber experiment. XX collected and analyzed the data and wrote a first draft of the manuscript. KSA, TD, SG, MH, MH, RH, RJ, RK, JVMS, EAS, YBS, BW, and SKW contributed to conducting the experiment. AS led the phytohormone analyses and acquired funding. CM led in conceiving the study, developing the experimental design, editing of the manuscript, and acquired funding. All authors reviewed the manuscript and gave final approval for publication.

### Supplementary Information

The online version contains supplementary material.

## Acknowledgements

We thank Ulrich Junghans, Petra Seibel, Ina Zimmer, and the EUS team for their help in the laboratory and rearing of plants; Tanja Bloss for the administration work and material preparation; Gunda Teichmann, Liv Kraack, and Frauke Harmel for rearing *P. nigra* trees and *L. dispar* caterpillars; Sarah Isabel Richard for preparing *Spodoptera exigua*; Claudia Schilke, Stefan Heiske, and Iris Klaiber for phytohormone extraction and running the analysis; Tanja Brendel, Felix Kreuzmann, Sunajana Chandra, Senén López Pérez, Joshua Flemming, and Hanna Greßmann for their great help in plant harvest.

