## Supplementary_Tables_Figures for "Common, species-specific, and accession-specific responses of foliar phytohormones and morphological traits to drought and herbivory"

**Table S1.** Time schedule of experiments

| | ***Tanacetum vulgare*** | ***Solanum dulcamara*** | ***Populus nigra*** |
|---|---|---|---|
| Plant cuttings preparation | 18/09/2024 | 27/01/2025 | 12/10/2024 |
| Transport plants to Munich | 18/09/2024 | 17/02/2025 | 16/12/2024 |
| Transfer plants into climate chambers | 04/12/2024 | 03/03/2025 | 16/06/2025 |
| [1]EK1+2: onset of drought treatment (defined as the day after the last uniform watering (day 0)) | 13/01/2025 | 11/03/2025 | 23/06/2025 |
| [2]EK3+4: onset of drought treatment | 14/01/2025 | 12/03/2025 | 24/06/2025 |
| Set larvae on leaf mix for adaptation | 21/01/2025 | 18/03/2025 | 30/06/2025 |
| EK1+2: Herbivory treatment | 23/01/2025 | 20/03/2025 | 04/07/2025 |
| EK3+4: Herbivory treatment | 24/01/2025 | 21/03/2025 | 05/07/2025 |
| EK 1+2: VOC collection | 26/01/2025 | 23/03/2025 | 07/07/2025 |
| EK 3+4: VOC collection | 27/01/2025 | 24/03/2025 | 08/07/2025 |
| EK1+2: Herbivore removal, shoot harvest, phenotyping | 27/01/2025 | 24/03/2025 | 08/07/2025 |
| EK3+4: Herbivore removal, shoot harvest, phenotyping | 28/01/2025 | 25/03/2025 | 09/07/2025 |
| EK1+2: Root harvest | 29/01/2025 | 26/03/2025 | 08/07/2025 |
| EK3+4: Root harvest | 30/01/2025 | 27/03/2025 | 09/07/2025 |

[1]EK1+2: main chambers one and two; [2]EK3+4: main chambers three and four

**Table S2.** Origin of accessions of *Tanacetum vulgare* individuals and their chemotypes

| Accession | Chemotype[1] | Coordinates (Country) | Collection date in field (collector) | Voucher ID |
|---|---|---|---|---|
| M6 | α-chrysanthenyl acetate (6_22) | 51°58.913"N 8°27.629"E (Germany) | 29/01/2019 (Elisabeth J. Eilers, Ruth Jakobs) | TBA[2] |
| | β-carvylacetate (6_27) | | | TBA |
| M9 | α-thujone (9_19) | 51°58.662"N 8°27.273" E(Germany) | 29/01/2019 (Elisabeth J. Eilers, Ruth Jakobs) | TBA |
| | α-thujone, β-thujone (9_35) | | | TBA |
| | artemisia ketone (9_77) | | | TBA |
| M11 | β-thujone (11_7) | 51°58.648"N 8°27.243"E (Germany) | 29/01/2019 (Elisabeth J. Eilers, Ruth Jakobs) | TBA |
| M16 | artemisia ketone, artemisyl acetate (16_40) | 51°58.601"N 8°27.173"E (Germany) | 29/01/2019 (Elisabeth J. Eilers, Ruth Jakobs) | TBA |
| | α-thujone, β-thujone (16_175) | | | TBA |
| M18 | camphene, cis-verbenol acetate (18_11) | 51°59.031"N 8°28.302"E (Germany) | 29/01/2019 (Elisabeth J. Eilers, Ruth Jakobs) | TBA |
| | (*Z*)-myroxide, santolina triene, artemisyl acetate (18_81) | | | TBA |
| M22 | artemisia ketone (22_5) | 51°58.994"N 8°28.388"E (Germany) | 29/01/2019 (Elisabeth J. Eilers, Ruth Jakobs) | TBA |
| | (*Z*)-myroxide, santolina triene, artemisyl acetate (22_16) | | | TBA |

Per accession (maternal origin) and offspring chemotype combination, 16 clones were produced from the offspring.

[1]The dominant leaf terpenoid(s) is (are) given. Chemotypes were determined in the seedling stage of the offspring plants

[2]TBA – to be added. The vouchers are currently prepared and voucher numbers will be added.

**Table S3.** Origin of accessions of *Solanum dulcamara* individuals

| Accession | Coordinates (Country) | Collection date in field (collector) | Selfed offspring generation (yes/no) | Voucher ID |
|---|---|---|---|---|
| SIE/A27 | 52°16'53.6"N 13°11'17.4"E (Germany) | 13/10/2013 (Tobias Lortzing) | yes | HOH-023157 |
| EBB/16 | 48°50'52.1"N 9°35'48.8"E (Germany) | 22/10/2020 (Anke Stepp-uhn) | no | HOH-023155 |
| EBB/18 | 48°51'11.3"N 9°35'36.9"E (Germany) | 22/06/2021 (Kruthika S Aragam) | no | HOH-023156 |
| OW09 | 51°51'36.3"N 5°54'03.5"E (Netherlands) | Autumn 2012 and 2013 (Qian Zhang, et al.) | no | TBA[1] |
| VW08 | 51°50'58.6"N 4°04'34.3"E (Netherlands) | Autumn 2012 and 2013 (Qian Zhang, et al.) | no | TBA |
| TW12 | 53°07'17.5"N 4°47'13.6"E (Netherlands) | Autumn 2012 and 2013 (Qian Zhang, et al.) | no | TBA |

Per accession, 32 clones were produced.

[1]TBA – to be added. The vouchers are currently prepared and voucher numbers will be added.

**Table S4.** Origin of accessions of *Populus nigra* individuals

| Accession | Sex | **Coordinates (Germany)** Topographic map sheet number - Municipality code – object # in this area; City of Marbach am Neckar[1] | Collection date in field (collector) | Voucher ID |
|---|---|---|---|---|
| R14 | Female | 7221-116- 01 | 2012-2013 by „Arbeitskreis zur Erhaltung der Neckarschwarzpappel, Stiftung Energie & Klimaschutz Badenn-Württemberg“[1], | TBA[2] |
| R19 | Female | 7421-116- 06 | | TBA |
| R20 | Female | 7421-116- 07 | | TBA |
| R25 | Male | 7019-118- 01 | | TBA |
| R30 | Male | 7021-118- 04 | | TBA |
| R32 | Male | 7021-118- 11 | | TBA |

Per accession, 16 clones were produced.

[1]https://www.energie-klimaschutz.de/wp-content/uploads/2015/01/Neckarschwarzpappel-Projektinformation-II.pdf

[2]TBA – to be added. The vouchers are currently prepared and voucher numbers will be added.

**Table S5A.** Watering regime (mL water/pot) for *Tanacetum vulgare* in the different treatment groups. The onset of drought was defined as the first day after the last uniform watering (13/01/2025 = day 0 for EK1 and EK2, 14.01.2025 = day 0 for EK3 and EK4). When two dates are listed, the first refers to watering in EK1 and EK2 and the second to watering in EK3 and EK4.

| | | Treatment groups | | | |
|---|---|---|---|---|---|
| **Date** | **Days of drought** | **Control** | **Herbivory** | **Drought** | **Combined** |
| 06/01/2025 | day-7 / day-8 | 200 | 200 | 200 | 200 |
| 09/01/2025* | day -4 / day-5 | 200 | 200 | 200 | 200 |
| 12/01/2025 | day -1 / day-2 | 200 | 200 | 200 | 200 |
| 15.,16/01/2025 | day +2 | 200 | 200 | 100[1] | 100[1] |
| 17.,18/01/2025 | day +4 | 200 | 200 | 100 | 100 |
| 19.,20/01/2025 | day +6 | 200 | 200 | 100 | 100 |
| 21.,22.01/2025 | day +8 | 200 | 200 | 100 | 100 |
| 23.,24.01/2025 | day +10 | 200 | 200 | 100 | 100 |
| 24.,26/01/2025 | day +12 | 200 | 200 | 100 | 100 |
| | **Total** | **1800** | **1800** | **1200** | **1200** |

*including 0.5% Hakaphos rot (Compo, GmbH, Germany) fertilizer

[1] 10 pots in drought and 13 pots in the combined treatment, respectively, received additional 50 mL to avoid wilting.

**Table S5B.** Watering regime (mL water/pot) for *Solanum dulcamara* in the different treatment groups. The onset of drought was defined as the first day after the last uniform watering (11/03/2025 = day 0 for EK1 and EK2, 12.03.2025 = day 0 for EK3 and EK4). When two dates are listed, the first refers to watering in EK1 and EK2 and the second to watering in EK3 and EK4.

| | | Treatment groups | | | |
|---|---|---|---|---|---|
| **Date** | **Days of drought** | **Control** | **Herbivory** | **Drought** | **Combined** |
| 06.,07/03/2025 | day -5 | 100 | 100 | 100 | 100 |
| 08.,09/03/2025 | day -3* | 100[1] | 100 | 100 | 100[1] |
| 10.,11/03/2025 | day -1 | 100 | 100 | 100 | 100 |
| 12.,13/03/2025 | day +1 | 100 | 100 | | |
| 14.,15/03/2025 | day +3 | 200 | 200 | 100 | 100 |
| 16.,17/03/2025 | day +5 | 200 | 200 | 100 | 100 |
| 18.,19/03/2025 | day +7 | 200 | 200 | 100[2] | 100[2] |
| 20.,21/03/2025 | day +9 | 300 | 300 | 150[2] | 150[2] |
| 22.,23/03/2025 | day + 11 | 300 | 300 | 150[2] | 150[2] |
| | **Total** | **1600** | **1600** | **900** | **900** |

[*]including 0.5% Hakaphos rot (Compo, GmbH, Germany) fertilizer

[1] Due to a malfunction in the automatic irrigation system, 12 plants in each of the control and combined treatment groups received 200 mL of water.

[2] 5 plants in the drought and 3 plants in the combined treatment received 50 mL less water due to their size.

**Table S5C.** Watering regime (mL water/pot) for *Populus nigra* in the different treatment groups. The onset of drought was defined as the first day after the last uniform watering (26.,27/06/2025 = day 0 for EK1 and EK2, 27/06/2025 = day 0 for EK3 and EK4). When two dates are listed, the first refers to watering in EK1 and EK2 and the second to watering in EK3 and EK4.

| | | Treatment groups | | | |
|---|---|---|---|---|---|
| **Date** | **Days of drought** | **Control** | **Herbivory** | **Drought** | **Combined** |
| 18/06/2025 | day -9 | 200 | 200 | 200 | 200 |
| 20/06/2025 | day -7 | 200 | 200 | 200 | 200 |
| 23/06/2025 | day -4 | 200 | 200 | 200 | 200 |
| 26.,27/06/2025 | day 0 | 200 | 200 | 100 | 100 |
| 29.,30/06/2025 | day +3 | 200 | 200 | 100 | 100 |
| 01.,02/07/2025 | day +5 | 200 | 200 | 100 | 100 |
| 03.,04/07/2025 | day +7 | 300 | 300 | 150 | 150 |
| | **Total** | **1500** | **1500** | **1050** | **1050** |

**Table S6A:** Soil water content (SWC, vol-%) in *Tanacetum vulgare* pots, determined using a soil moisture meter (HH2) connected to a ThetaProbe (ML2x; Delta-T Devices. EKs denote the experimental chambers in which measurements were performed. Values are presented as mean ± standard deviation (SD) and standard error (SE). When four chambers were measured, *n* = 48 per treatment; when two chambers were measured, *n* = 24 per treatment.

| Date | Day of drought | SWC | Treatment | | | | EKs |
|---|---|---|---|---|---|---|---|
| | | | Control | Herbivory | Drought | Combined | |
| | | | (vol-%) | (vol-%) | (vol-%) | (vol-%) | |
| 07/01/2025 | day -6 | mean | 28.3 | 26.9 | 26.9 | 25.9 | EK 1-4 |
| | | SD | 11.2 | 10.3 | 11.0 | 9.7 | |
| | | SE | 1.7 | 1.6 | 1.7 | 1.5 | |
| 13/01/2025 | day 0 | mean | 25.9 | 26.6 | 25.1 | 25.8 | EK 1-4 |
| | | SD | 14.0 | 11.7 | 11.1 | 10.6 | |
| | | SE | 2.0 | 1.7 | 1.6 | 1.5 | |
| 16/01/2025 | day + 3 | mean | | | 11.9 | 12.5 | EK 1-4 |
| | | SD | | | 5.7 | 6.8 | |
| | | SE | | | 0.8 | 1.0 | |
| 17/01/2025 | day +4 | mean | | | 9.2 | 9.6 | EK 3+4 |
| | | SD | | | 4.7 | 5.8 | |
| | | SE | | | 1.0 | 1.2 | |
| 20/01/2025 | day +7 | mean | 30.4 | 27.9 | 11.6 | 12.6 | EK 1+2 |
| | | SD | 14.2 | 10.2 | 4.0 | 5.8 | |
| | | SE | 2.9 | 2.1 | 0.8 | 1.2 | |
| 21/01/2025 | day +8 | mean | 26.3 | 27.7 | 13.2 | 12.5 | EK 3+4 |
| | | SD | 11.8 | 12.8 | 8.5 | 8.3 | |
| | | SE | 2.4 | 2.6 | 1.7 | 1.7 | |
| 22/01/2025 | day +9 | mean | | | 11.1 | 11.3 | EK 1+2 |
| | | SD | | | 5.1 | 5.5 | |
| | | SE | | | 1.0 | 1.1 | |
| 23/01/2025 | day +10 | mean | | | 11.7 | 11.2 | EK 3+4 |
| | | SD | | | 6.3 | 6.6 | |
| | | SE | | | 1.3 | 1.3 | |

**Table S6B:** Soil water content (SWC, vol-%) in *Solanum dulcamara* pots, determined using a soil moisture meter (HH2) connected to a ThetaProbe (ML2x; Delta-T Devices). Values are presented as mean ± standard deviation (SD) and standard error (SE), *n* = 48 per treatment. EKs denote the experimental chambers in which measurements were performed. When two dates are listed, the first refers to watering in EK1 and EK2 and the second to watering in EK3 and EK4.

| Date | Day of drought | SWC | Treatment | | | | EKs |
|---|---|---|---|---|---|---|---|
| | | | Control | Herbivory | Drought | Combined | |
| | | | vol-% | vol-% | vol-% | vol-% | |
| 06.,07/03/2025 | day -5 | mean | 39.2 | 39.1 | 38.0 | 39.1 | EK 1-4 |
| | | SD | 5.8 | 5.5 | 4.9 | 6.6 | |
| | | SE | 0.8 | 0.8 | 0.7 | 1.0 | |
| 11.,12/03/2025 | day 0 | mean | 30.2 | 31.8 | 30.0 | 32.0 | EK 1-4 |
| | | SD | 8.1 | 8.8 | 8.7 | 8.6 | |
| | | SE | 1.2 | 1.3 | 1.3 | 1.2 | |
| 13.,14/03/2025 | day +2 | mean | 33.6 | 34.5 | 22.4 | 24.3 | EK 1-4 |
| | | SD | 7.6 | 8.9 | 9.4 | 8.9 | |
| | | SE | 1.1 | 1.3 | 1.4 | 1.3 | |
| 17.,18/03/2025 | day +6 | mean | 35.8 | 35.4 | 19.2 | 19.5 | EK 1-4 |
| | | SD | 8.1 | 8.9 | 9.9 | 9.5 | |
| | | SE | 1.2 | 1.3 | 1.4 | 1.4 | |
| 19.,20/03/2025 | day +8 | mean | 35.9 | 35.6 | 18.6 | 18.0 | EK 1-4 |
| | | SD | 7.9 | 8.5 | 10.2 | 8.8 | |
| | | SE | 1.1 | 1.2 | 1.5 | 1.3 | |

**Table S6C**: Soil water content (SWC, vol-%) in *Populus nigra* pots, determined using a soil moisture meter (HH2) connected to a ThetaProbe (ML2x; Delta-T Devices). Values are presented as mean ± standard deviation (SD) and standard error (SE), *n* = 48 per treatment. EKs denote the experimental chambers in which measurements were performed. When two dates are listed, the first refers to watering in EK1 and EK2 and the second to watering in EK3 and EK4.

| Date | Day of drought | SWC | Treatment | | | | EKS |
|---|---|---|---|---|---|---|---|
| | | | **Control** | **Herbivory** | **Drought** | **Combined** | |
| | | | vol-% | vol-% | vol-% | vol-% | |
| 22.,23/06/2025 | day -5 | mean | 23.9 | 18.9 | 19.6 | 20.6 | EK 1-4 |
| | | SD | 12.9 | 7.6 | 7.8 | 8.9 | |
| | | SE | 1.9 | 1.1 | 1.1 | 1.3 | |
| 26.,27/06/2025 | day 0 | mean | 30.6 | 24.7 | 18.9 | 21.1 | EK 1-4 |
| | | SD | 11.0 | 8.6 | 6.7 | 9.7 | |
| | | SE | 1.6 | 1.2 | 1.0 | 1.4 | |
| 02.,03/07/2025 | day +6 | mean | 27.1 | 23.8 | 12.3 | 13.9 | EK 1-4 |
| | | SD | 12.2 | 8.8 | 4.3 | 7.1 | |
| | | SE | 1.8 | 1.3 | 0.6 | 1.0 | |
| 07.,08/07/2025 | day +10 | mean | 26.6 | 21.5 | 9.1 | 10.2 | EK 1-4 |
| | | SD | 11.7 | 9.5 | 2.5 | 6.0 | |
| | | SE | 1.7 | 1.4 | 0.4 | 0.9 | |

**Table S7.** Means (standard deviations) of phenotypic plasticity (measured as RDPI[1]) of traits of *Populus nigra* of different sex, exposed to different challenges (combined: drought and herbivory).

| Trait | Treatment | Female<br>df = 12 | Male<br>df = 12 | |
|---|---|---|---|---|
| JA concentration | Herbivory | 0.656 (0.071) | 0.705 (0.053) | |
| | Drought | 0.323 (0.052) | 0.300 (0.063) | |
| | Combined | 0.479 (0.083) | 0.510 (0.073) | |
| JA-Ile concentration | Herbivory | 0.487 (0.102) | 0.558 (0.075) | |
| | Drought | 0.372 (0.086) | 0.513 (0.079) | |
| | Combined | 0.337 (0.068) | 0.493 (0.061) | |
| SA concentration | Herbivory | 0.266 (0.044) | 0.161 (0.035) | Sex: |
| | Drought | 0.367 (0.061) | 0.193 (0.037) | $\chi^2 = 5.57$, |
| | Combined | 0.272 (0.049) | 0.254 (0.061) | $p = 0.02$ |
| ABA concentration | Herbivory | 0.406 (0.057) | 0.411 (0.077) | |
| | Drought | 0.856 (0.055) | 0.797 (0.057) | |
| | Combined | 0.819 (0.064) | 0.822 (0.042) | |
| IAA concentration | Herbivory | 0.189 (0.045) | 0.190 (0.033) | |
| | Drought | 0.276 (0.049) | 0.149 (0.031) | |
| | Combined | 0.263 (0.059) | 0.166 (0.032) | |
| Height | Herbivory | 0.047 (0.013) | 0.056 (0.011) | |
| | Drought | 0.056 (0.014) | 0.093 (0.016) | |
| | Combined | 0.067 (0.011) | 0.065 (0.015) | |
| Leaf number | Herbivory | 0.057 (0.019) | 0.067 (0.012) | |
| | Drought | 0.085 (0.025) | 0.064 (0.016) | |
| | Combined | 0.094 (0.030) | 0.102 (0.018) | |
| Total leaf dry mass | Herbivory | 0.091 (0.023) | 0.069 (0.023) | |
| | Drought | 0.128 (0.026) | 0.103 (0.026) | |
| | Combined | 0.122 (0.018) | 0.089 (0.020) | |
| Total stem dry mass | Herbivory | 0.136 (0.028) | 0.091 (0.023) | |
| | Drought | 0.092 (0.016) | 0.136 (0.036) | |
| | Combined | 0.113 (0.012) | 0.111 (0.026) | |
| Total aboveground dry mass | Herbivory | 0.104 (0.021) | 0.068 (0.021) | |
| | Drought | 0.096 (0.019) | 0.097 (0.031) | |
| | Combined | 0.095 (0.016) | 0.061 (0.020) | |
| Root dry mass | Herbivory | 0.104 (0.020) | 0.098 (0.024) | |
| | Drought | 0.119 (0.023) | 0.150 (0.035) | |
| | Combined | 0.093 (0.022) | 0.160 (0.025) | |
| Total dry mass | Herbivory | 0.092 (0.018) | 0.077 (0.021) | |
| | Drought | 0.085 (0.017) | 0.113 (0.033) | |
| | Combined | 0.077 (0.016) | 0.087 (0.021) | |
| Root shoot ratio | Herbivory | 0.090 (0.023) | 0.059 (0.015) | |
| | Drought | 0.104 (0.032) | 0.073 (0.017) | |
| | Combined | 0.103 (0.024) | 0.128 (0.026) | |
| Leaf water content | Herbivory | 0.016 (0.003) | 0.031 (0.010) | Sex: |
| | Drought | 0.026 (0.006) | 0.029 (0.005) | $\chi^2 = 3.59$, |
| | Combined | 0.036 (0.008) | 0.059 (0.018) | $p = 0.058$ |
| Stem specific density | Herbivory | 0.165 (0.028) | 0.216 (0.047) | |
| | Drought | 0.178 (0.033) | 0.209 (0.041) | |
| | Combined | 0.157 (0.043) | 0.108 (0.028) | |

[1]RDPI: $(|x_c - x_s|)/(|x_c + x_s|)$ where $X_c$ and $X_s$ represent the different trait values of clones kept under control (c) and challenge (s) conditions.

**Table S8: Global goodness-of-fit of each structure equation model (SEM)**

| Species | SEM | Fisher' C | *p* value | *df* |
|---|---|---|---|---|
| *Tanacetume vulgare* | Treatment and accession | 59.61 | *0.09* | 46 |
| *Solanum dulcamara* | Treatment and accession | 70.76 | *0.09* | 56 |
| *Populus nigra* | Treatment and genotype | 80.05 | 0.66 | 86 |
| | Treatment and sex | 91.14 | **0.01** | 66 |

Significant *p*-values of ($p \leq 0.05$) are highlighted in bold, marginally significant ($p < 0.1$) in italics.

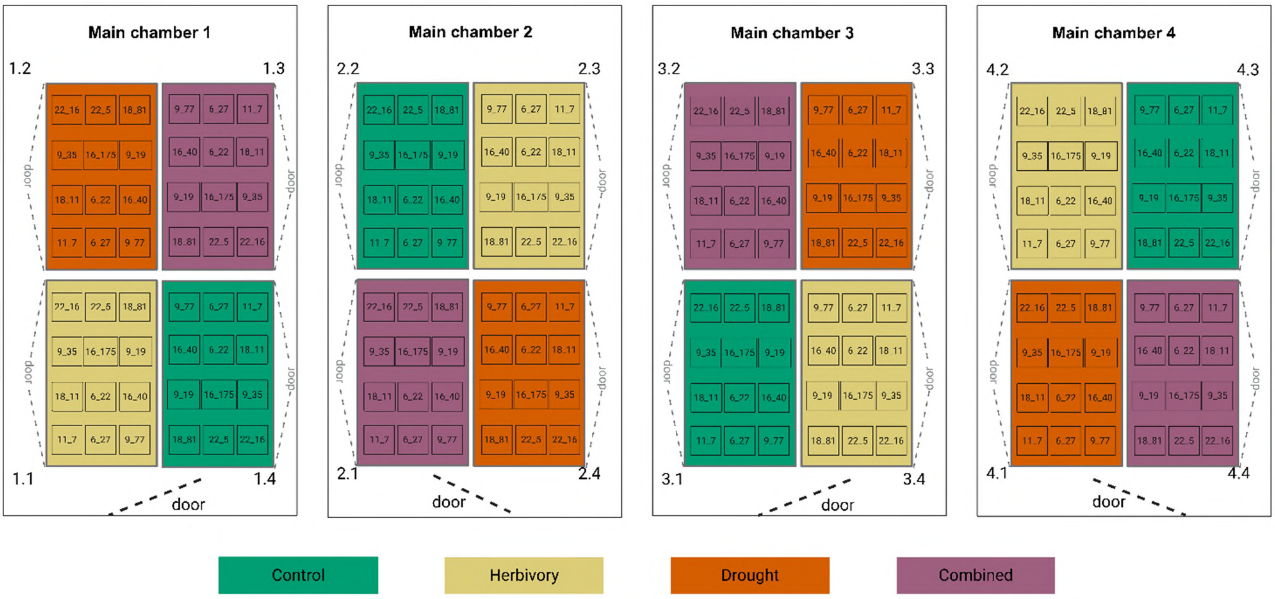


**Figure S1.** Set-up of *Tanacetum vulgare* clones in experimental chambers. The experimental set-up consisted of four main chambers, with four sub-chambers in each main chamber. Clonal plants experienced one of the four treatments control, drought, insect herbivory and combined challenges (drought and herbivory) in one of four sub-chambers within the main chambers. Clones of one plant individual are indicated by an ID consisting of the maternal origin and the offspring number (for codes see Table S2).

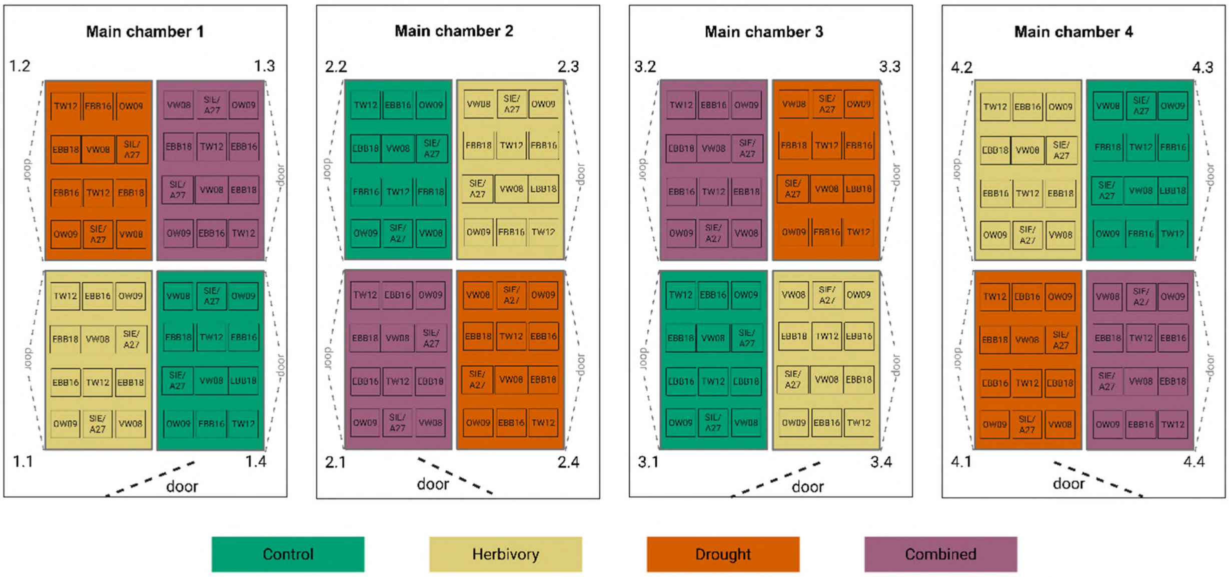


**Figure S2.** Set-up of *Solanum dulcamara* plants in experimental chambers. The experimental set-up consisted of four main chambers, with four sub-chambers in each main chamber. Clonal plants experienced one of the four treatments control, drought, insect herbivory and combined challenges (drought and herbivory) in one of four sub-chambers within the main chambers. Clones of one plant individual are indicated by an ID (for codes see Table S3).

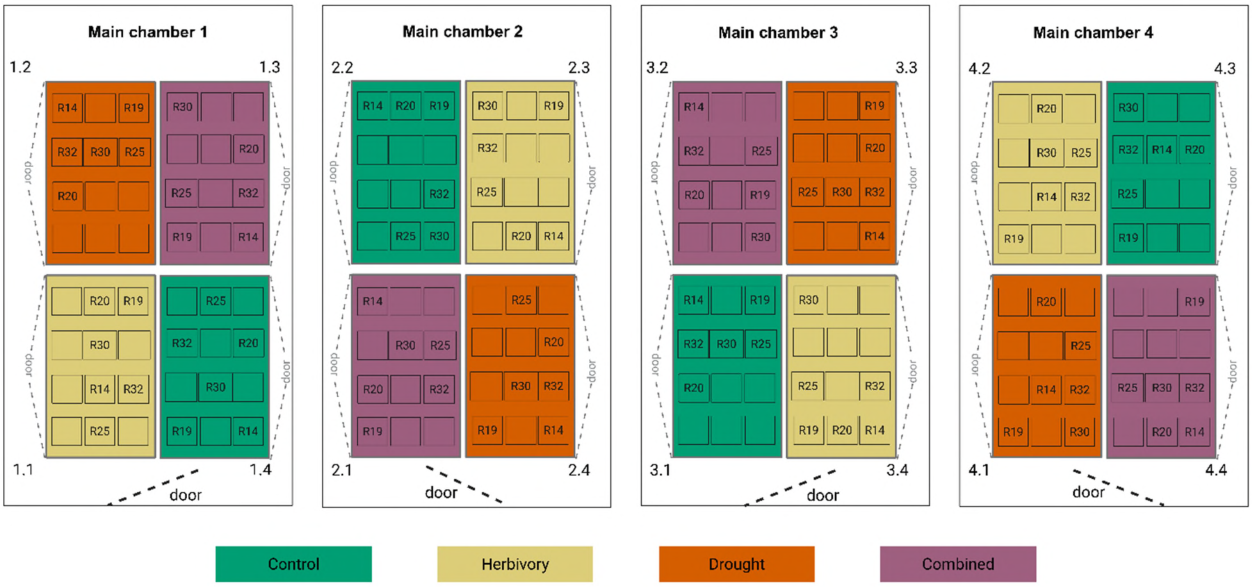


**Figure S3.** Set-up of *Populus nigra* plants in experimental chambers. The experimental set-up consisted of four main chambers, with four sub-chambers in each main chamber. Clonal plants experienced one of the four treatments control, drought, insect herbivory and combined challenges (drought and herbivory) in one of four sub-chambers within the main chambers. Clones of one plant individual are indicated by an ID (for codes see Table S4).

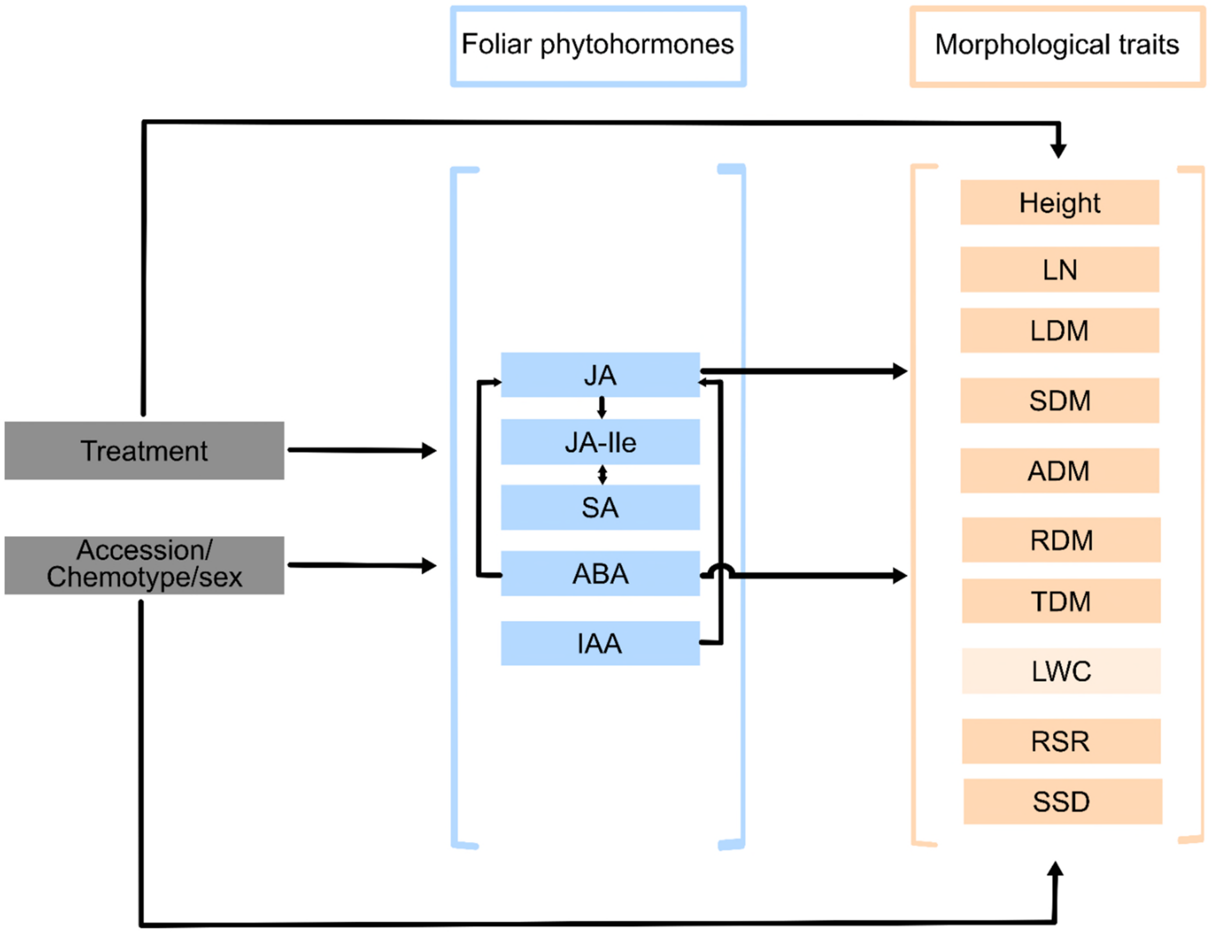


**Figure S4.** Graphical representation of the hypothesized causal flow of how foliar phytohormones regulated morphological responses to treatments and accession/chemotype/sex. Phytohormones (blue boxes) mediate the effect of treatments and accession/chemotype/sex (grey boxes) on morphological traits (peach boxes). The phytohormones are JA: jasmonic acid; JA-Ile: jasmonyl isoleucine; SA: salicylic acid; ABA: abscisic acid; IAA: indole-3-acetic acid. The morphological traits are: LN: leaf number; LDM: leaf dry mass; SDM: stem dry mass; ADM: aboveground dry mass; RDM: root dry mass; TDM: total dry mass; LWC: leaf water content; RSR: root-shoot ratio; SSD: specific stem density. LWC in light peach box is only affected by treatment and accession /sex.

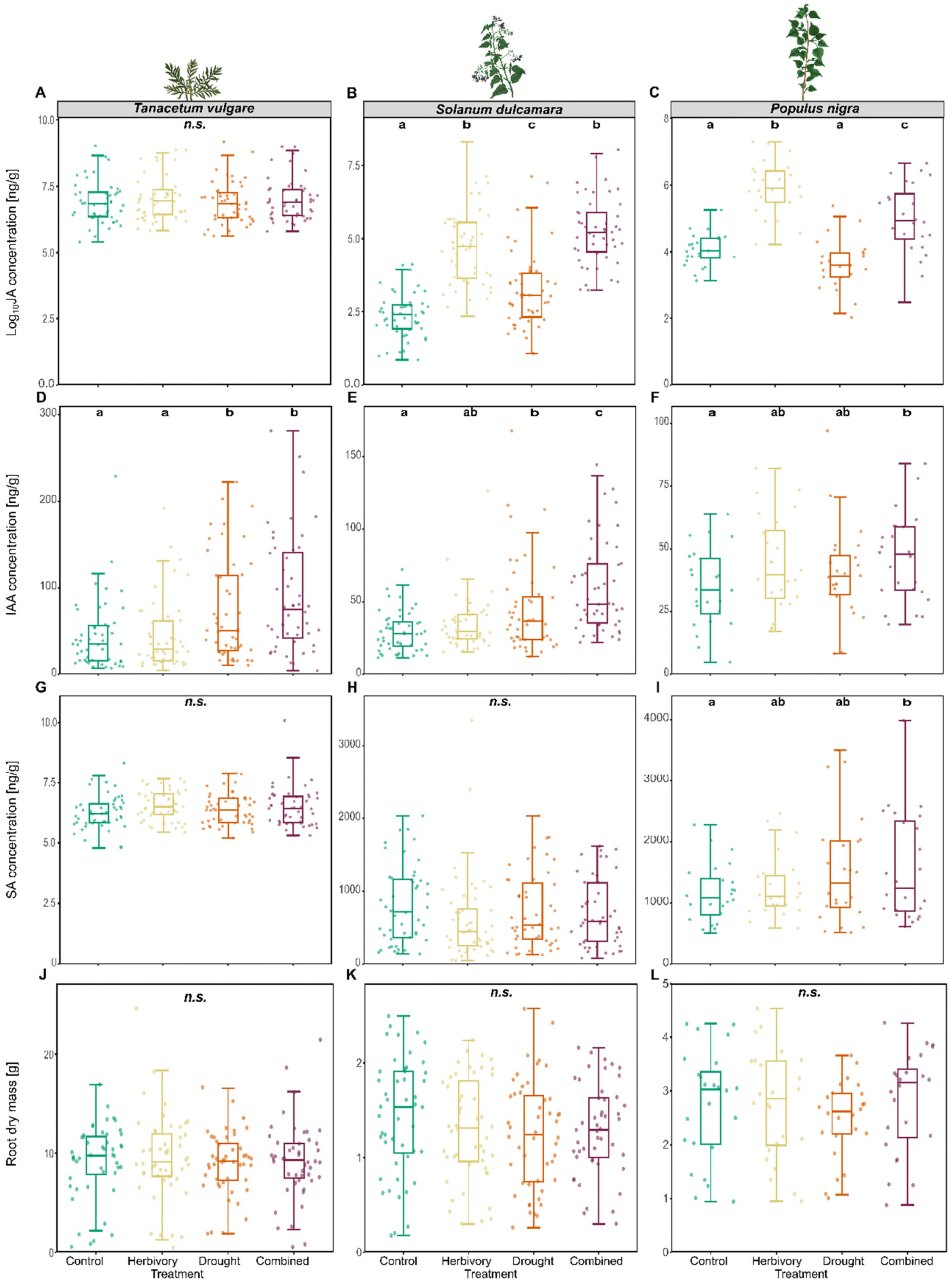


**Figure S5.** Boxplots of foliar phytohormone concentrations (JA: jasmonic acid; IAA: indole acetate acid; SA: salicylic acid) and root dry mass as morphological trait across treatments (combined: herbivory and drought) of each species, *Tanacetum vulgare* (A, D, G, J), *Solanum dulcamara* (B, E, H, K), and *Populus nigra* (C, F, I, L). Data are presented as boxplots, with medians, interquartile ranges (IQR, boxes), and whiskers extending to the most extreme values with max. 1.5 times the IQR. Individual values are plotted as dots; *n* = 48 per treatment for *T. vulgare* and *S. dulcamara*, n = 24 per treatment for *P. nigra*. Different letters indicate statistically significant differences (F-test, $p < 0.05$); *n.s.*: not significant. Please note the different scaling on the y-axes.

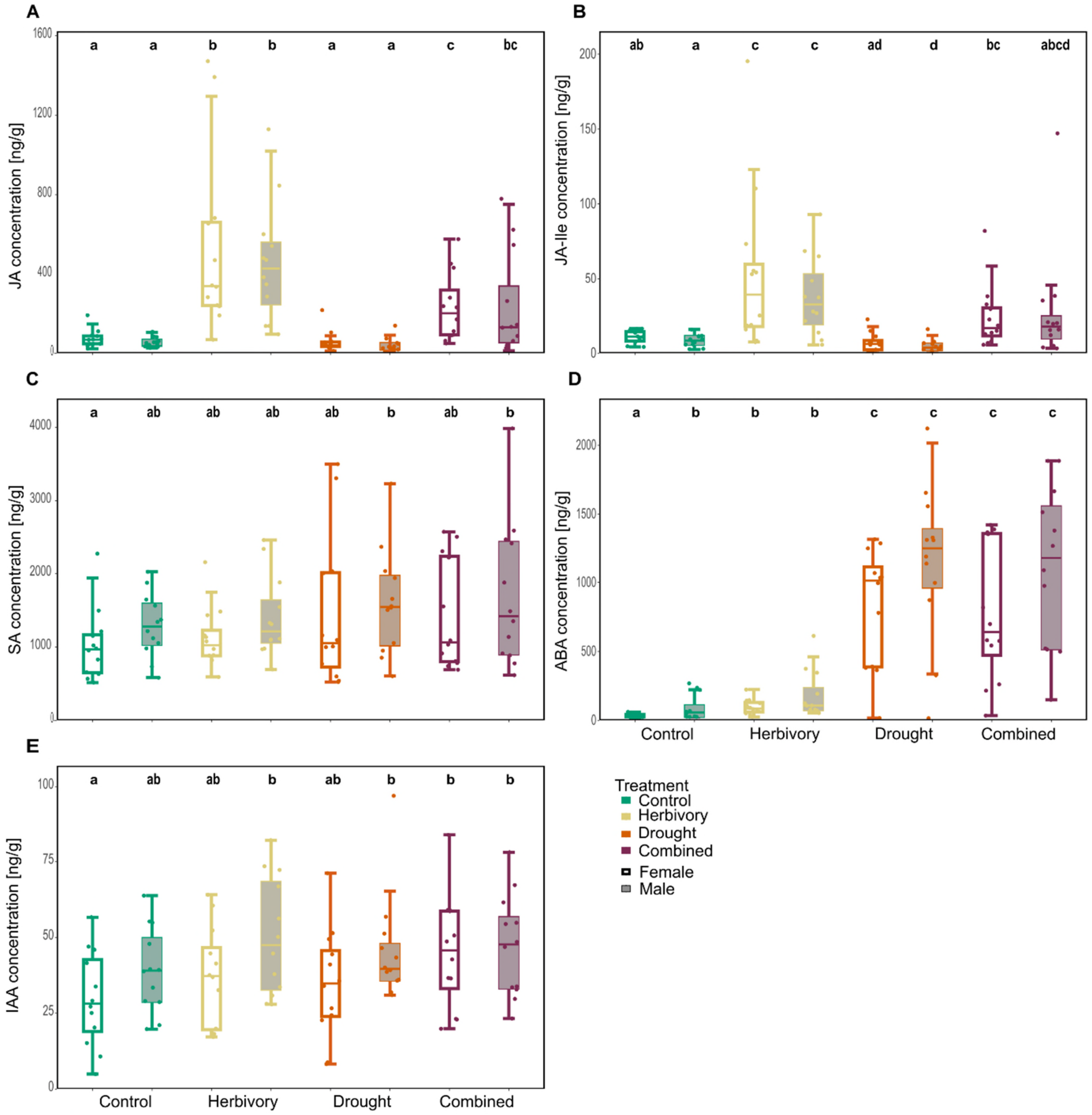


**Figure S6.** Boxplots of foliar phytohormone concentrations [JA: jasmonic acid (A); JA-Ile: jasmononyl isoleucine (B); SA: salicylic acid (C); ABA: abscisic acid (D); IAA: indole acetate acid (E)) across treatments (combined: herbivory and drought) and sex of *Populus nigra*. Data are presented as boxplots, with medians, interquartile ranges (IQR, boxes), and whiskers extending to the most extreme values with max. 1.5 times the IQR. Individual values are plotted as dots; n = 24 per treatment for *P. nigra*. Different letters indicate statistically significant differences (F-test, $p < 0.05$).

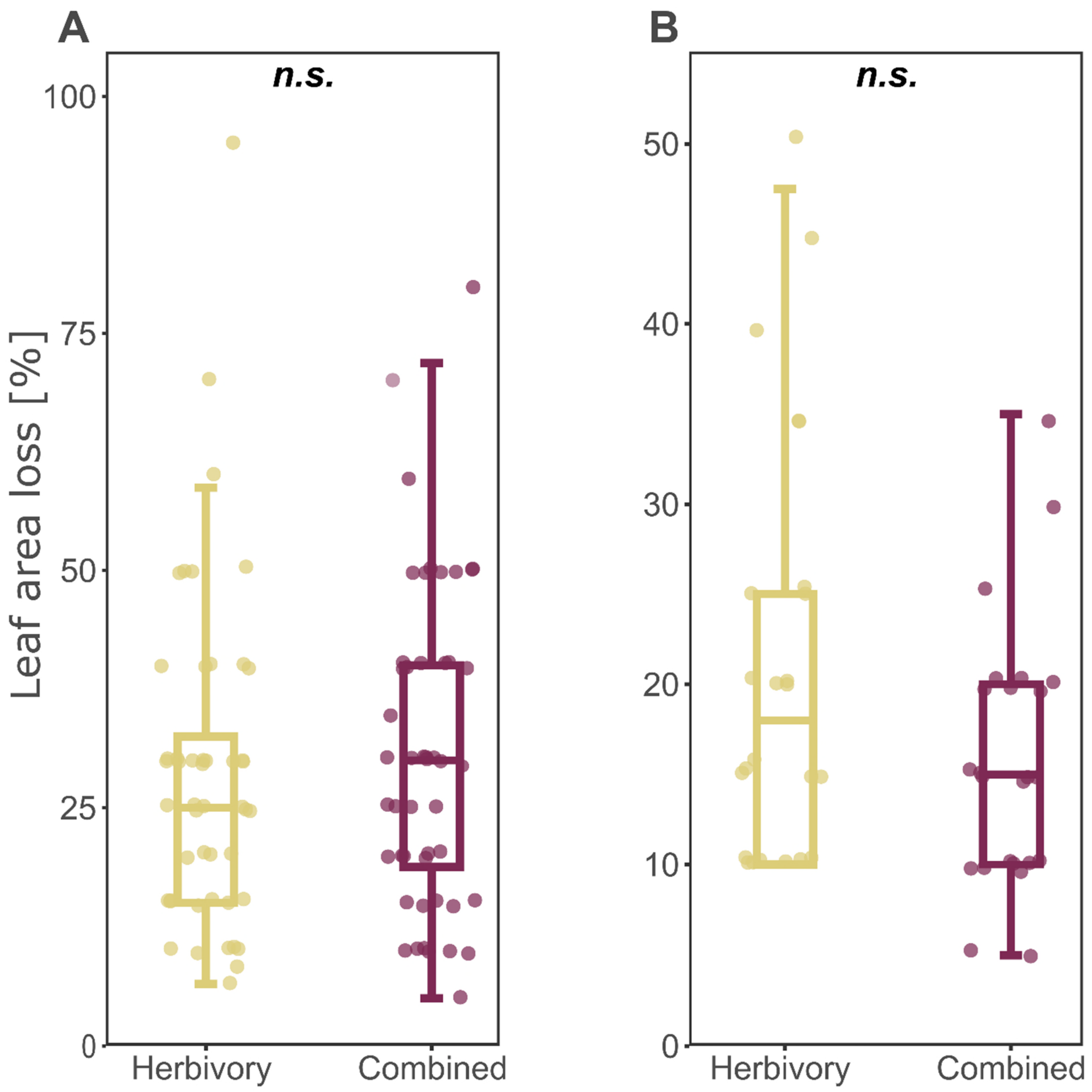


**Figure S7.** Boxplots of estimated leaf area loss in *Solanum dulcamara* (A), and *Populus nigra* (B) under herbivory or a combined (herbivory and drought) treatment. Observers were trained using ZAX herbivory images until they reached at least 10% estimation accuracy prior to the experiment. Leaf area loss was not assessed for *Tanacetum vulgare* because larvae of *Spodoptera exigua* primarily fed on the petiole rather than the leaf blade. Data are presented as boxplots, with medians, interquartile ranges (IQR, boxes), and whiskers extending to the most extreme values with max. 1.5 times the IQR. Individual values are plotted as dots; $n$ = 48 per treatment for *S. dulcamara*, n = 24 per treatment for *P. nigra*. No significant differences were found (*n.s.*, Mann-Whitney U test test, $p > 0.05$).

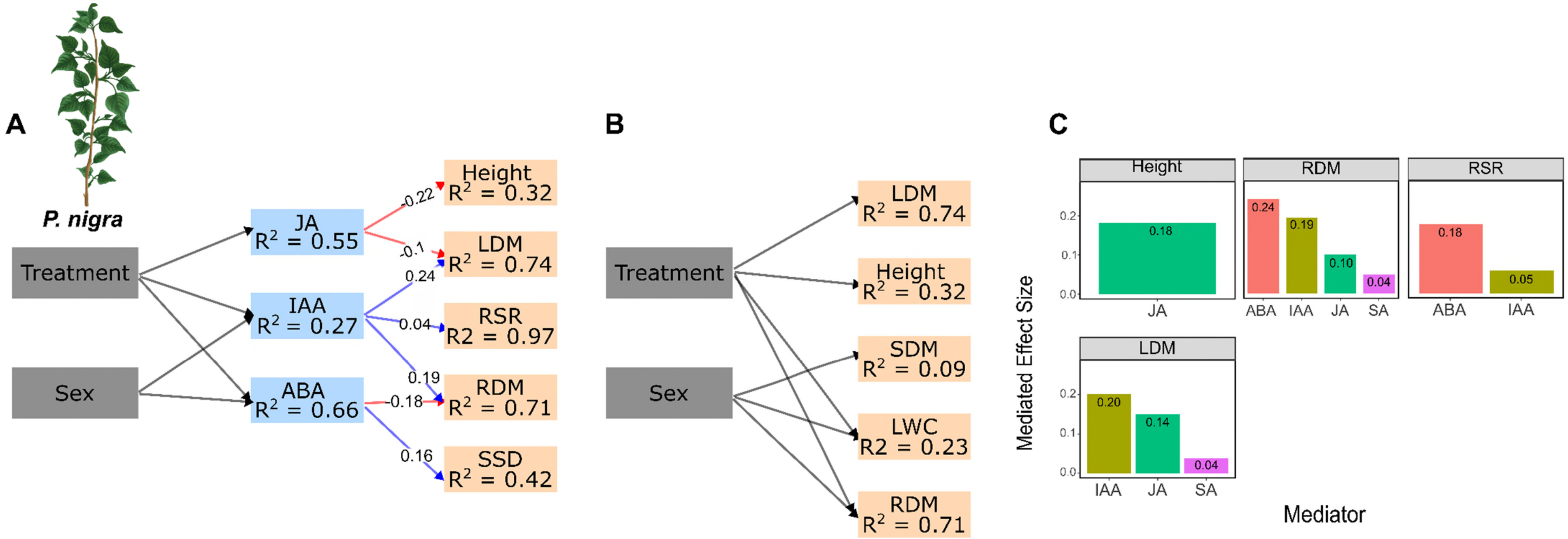


**Figure S8.** Structural equation models (SEMs) showing how foliar phytohormones regulated morphological responses to treatments in *Populus nigra*. Direct relationships between treatment and sex on foliar phytohormone concentrations, and between phytohormones and morphological traits in *P. nigra* (A). Numbers on each arrow represent standardized effect sizes. Blue arrows indicate positive relationships; red arrows indicate negative ones. Direct effects of treatment and sex on morphological traits in *Populus nigra* (B). The SEM showed imperfect fits, because some paths without biological causes were intentionally excluded (*Populus nigra*: Fisher's C = 94.14, *df* = 66, *p* = 0.01). Coefficients of determination ($R^2$) for each endogenous variable are shown in the corresponding boxes. For each SEM, the magnitudes of phytohormone mediator effects (i.e., all indirect pathways linking treatment and sex to morphological traits) are shown as bar plots (C: *Populus nigra*). Bootstrapped mediator effects are presented as absolute values to allow comparison of effect strength regardless of direction. The morphological traits are: LN: leaf number; LDM: leaf dry mass; SDM: stem dry mass; RDM: root dry mass; RSR: root-shoot ratio; LWC: leaf water content; SSD: specific stem density.
